# Transcription factors form a ternary complex with NIPBL/MAU2 to localize cohesin at enhancers

**DOI:** 10.1101/2024.12.09.627537

**Authors:** Gregory Fettweis, Kaustubh Wagh, Diana A. Stavreva, Alba Jiménez-Panizo, Sohyoung Kim, Michelle Lion, Andrea Alegre-Martí, Lorenzo Rinaldi, Thomas A. Johnson, Manan Krishnamurthy, Li Wang, David A. Ball, Tatiana S. Karpova, Arpita Upadhyaya, Didier Vertommen, Juan Fernández Recio, Eva Estébanez-Perpiñá, Franck Dequiedt, Gordon L. Hager

## Abstract

While the cohesin complex is a key player in genome architecture, how it localizes to specific chromatin sites is not understood. Recently, we and others have proposed that direct interactions with transcription factors lead to the localization of the cohesin-loader complex (NIPBL/MAU2) within enhancers. Here, we identify two clusters of LxxLL motifs within the NIPBL sequence that regulate NIPBL dynamics, interactome, and NIPBL-dependent transcriptional programs. One of these clusters interacts with MAU2 and is necessary for the maintenance of the NIPBL-MAU2 heterodimer. The second cluster binds specifically to the ligand-binding domains of steroid receptors. For the glucocorticoid receptor (GR), we examine in detail its interaction surfaces with NIPBL and MAU2. Using AlphaFold2 and molecular docking algorithms, we uncover a GR-NIPBL-MAU2 ternary complex and describe its importance in GR-dependent gene regulation. Finally, we show that multiple transcription factors interact with NIPBL-MAU2, likely using interfaces other than those characterized for GR.

## INTRODUCTION

Chromatin loops are at the base of 3D genome organization and are highly dynamic structures that are formed by cohesin complex-mediated extrusion (1). The ring-shaped cohesin assembly consists of SMC1, SMC3, RAD21, and either STAG1 or STAG2. The NIPBL/MAU2 heterodimer is widely believed to be the cohesin loader (2). Additionally, NIPBL stimulates the ATPase activity of the cohesin complex, promoting ATP-dependent cohesin translocation, resulting in loop extrusion (3). Loop enlargement continues until the cohesin complex either encounters a roadblock or is unloaded (4-6). Importantly, the cohesin complex is enriched at active enhancers and promoters, which could serve as cohesin loading sites (7-12). However, Banigan *et al*. showed that cohesin does not preferentially load at active transcription start sites and instead accumulates there due to obstruction by RNA Pol II (13). Taken together, these data suggest that enhancers are likely sites of cohesin loading. Given that NIPBL cannot recognize specific DNA sequences, how does it localize to specific enhancers (7, 14,15)?

We have previously proposed that transcription factors (TFs), which bind specific DNA sequences within enhancers and promoter-proximal regions, associate with NIPBL to localize cohesin to their target enhancers (7). Specifically, using the glucocorticoid receptor (GR), a hormone-inducible TF that belongs to the nuclear receptor (NR) superfamily, we showed that NIPBL and SMC1 bind to GR binding sites in a hor-mone-dependent manner. GR binding to enhancers triggers loop extrusion, and depletion of the cohesin subunit RAD21 impairs gene regulation by two ligand-inducible TFs (GR and NFκB) (7). Indeed, several TFs, chromatin remodelers, and other co-regulators have been shown to interact with NIPBL (7,16-21). However, the NIPBL/TF interaction surface(s) are still unknown.

In this study, we identify two clusters (named C1 and C2) of LxxLL motifs (where L stands for leucine and x can be any amino acid) within NIPBL that regulate its nuclear dynamics. LxxLL motifs are known to be enriched in TFs and facilitate protein-proteins interactions (PPIs) within transcription regulatory complexes (22,23). We show that NIPBL-C1 and C2 recruit a large variety of chromatin-associated proteins, including TFs. We further demonstrate that many TFs belonging to different superfamilies can interact directly with both NIPBL and MAU2, and that C1 is necessary to maintain the NIPBL-MAU2 heterodimer. Using surface plasmon resonance (SPR), we further show that several steroid receptors (SRs) interact with C2 through their ligand-binding domains (LBDs). AlphaFold and docking experiments predicted that NIPBL-C2 docks within the activation function-2 (AF-2) pocket of SRs. We combine these data with an Al-phaFold-generated model of the MAU2-C1 interaction to propose that TFs form a ternary complex with NIPBL/MAU2. Finally, we demonstrate the importance of these interactions in GR-mediated gene regulation.

## RESULTS

### Two Leu-rich clusters within NIPBL determine its nuclear dynamics and function

NIPBL is a large protein consisting of over 2700 residues with a mostly disordered N-terminal domain (NTD) consisting of approximately 1200 amino-acids (**Figure 1A, B**). NIPBL contains three evolutionary conserved Leucine (Leu)-rich clusters, each containing multiple LxxLL motifs (**Figure 1A, C**). Considering that NIPBL is enriched at TF binding sites (7-9), and can potentially directly interact with TFs (16,18,19), we set out to investigate the role of these three LxxLL motif clusters, referred to as C1, C2, and C3 in this manuscript, in promoting the interactions between NIPBL and other nuclear proteins.

**Figure 1:**
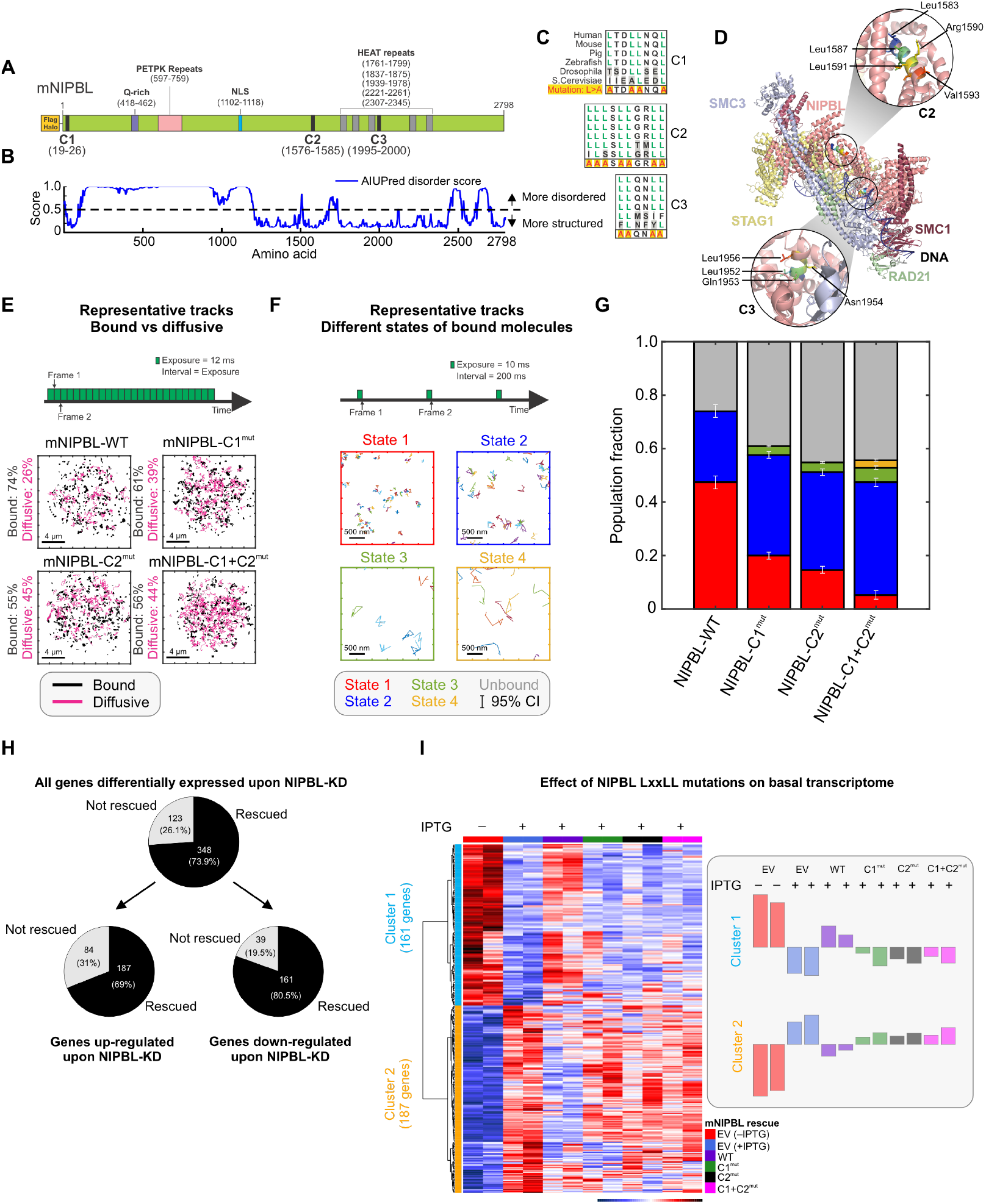
LxxLL mutations alter NIPBL dynamics and basal transcriptome. **(A)** Domain organization of *Mus musculus* NIPBL (mNIPBL) including the ectopic FLAG-Halo protein tags at the N-terminus of the protein. The three annotated LxxLL motif clusters are indicated as C1, C2, and C3. **(B)** AIUPred disorder score for mNIPBL. **(C)** Conservation of NIPBL LxxLL motif clusters across species. **(D)** Overall structure of the human cohesin-NIPBL-DNA complex solved by cryo-EM (24). The C2 and C3 clusters are indicated in the insets. **(E)** Fast SMT protocol (top) and 500 randomly selected single molecule trajectories for indicated species (bottom). Bound and diffusive fractions are indicated on the left of the respective panels. Scale bar is 4 μm. N_cells_/N_tracks_ = 70/15,394 (WT), 142/82,691 (C1^mut^), 76/46,879 (C2^mut^), 75/27,535 (C1+C2^mut^). **(F)** Intermediate SMT protocol captures the motion of bound NIPBL molecules (top). Representative trajectories of molecules in each of the four detected mobility states (bottom). Scale bar is 500 nm. N_cells_/N_tracks_/N_sub-tracks_ = 53/1,496/7,004 (WT), 65/5,769/19,632 (C1^mut^), 69/4,398/22,317 (C2^mut^), 67/1,746/7,729 (C1+C2^mut^). **(G)** Population fractions for indicated mNIPBL species. Error bars = 95% confidence interval. **(H)** Differentially expressed genes upon NIPBL-KD rescued by 3xFLAG-Halo-mNIPBL-WT expression under basal conditions. **(I)** (Left) Heatmap of the 348 genes rescued by ectopic 3xFLAG-Halo-NIPBL-WT compared across the cell lines. (Right) Average expression bar plots for the genes in each cluster. See also, Figure S1, Videos S1-S2.

Since the structure of the human NIPBL-cohesin-DNA complex has been recently solved by cryo-EM (24), we first asked whether the C2 (human aa1583-1592/mouse aa1576-1585) and C3 (human aa1951-1956/mouse aa1995-2000) clusters were topologically available for binding other proteins. C1 (human and mouse aa19-26) is in the disordered NTD of NIPBL (**Figure 1A, B**), and hence cannot be visualized in the cryo-EM structure. However, because it is embedded in a disordered region of NIPBL, it is likely available for interactions. In the cryo-EM structure, C2 is fully exposed while C3 is occluded by RAD21 and SMC3 (**Figure 1D**). Thus, in the rest of this study, we focused on C1 and C2, which are likely to be surface-exposed and accessible for PPIs.

To test the role of C1 and C2 in driving NIPBL-TF interactions, we built an IPTG-induced shRNA-based NIPBL knock-down/rescue system in 3134 mouse mammary adenocarcinoma cells (see Methods, Figure S1A), selecting a single-cell clone (cl15) that showed ∼70% knockdown (KD) of *Nipbl* by qPCR (**Figure S1A**). In the cl15 background, we then generated individual cell lines constitutively expressing 3xFLAG-Halo-mNIPBL-WT (cl15-WT), mutants harboring LxxLL mutations (Leu>Ala) in one or both of C1 and C2 clusters (cl15-C1^mut^, C2^mut^, and C1+C2^mut^, respectively), or 3xFLAG-Halo alone (empty-vector, cl15-EV) (**Figures 1C, S1A, see Methods**). As LxxLL motifs interact with partner proteins largely through hydrophobic interactions (22), replacing the non-polar hydrophobic Leu residues with smaller, less hydrophobic residues such as Alanine (Ala), is known to weaken LxxLL motif-driven interactions (25). RNA-seq reads show that the WT and C2^mut^ constructs are expressed to similar levels while the C1^mut^ and C1+C2^mut^ are expressed twice as much as the WT and C2^mut^ (**Figure S1B**).

Single-molecule tracking (SMT) is a powerful technology to investigate the spatiotemporal dynamics of proteins *in vivo* (26,27), and we used SMT to ask whether C1 and C2 regulate the intranuclear binding dynamics of NIPBL (**Figures 1E-G**). We first used a fast-imaging modality to track both diffusing and bound molecules (27) (**Figure 1E, Video S1**). Mutations in C1 and/or C2 resulted in a 13-19% decrease in the chromatin-bound fraction, indicating that both LxxLL clusters are important for the recruitment of NIPBL onto chromatin (**Figure 1E, G**).

Our previous work has shown that an intermediate SMT regime (short exposures but longer intervals) allows for the identification of two distinct low-mobility states for ‘bound’ molecules (28) (**Figure 1F, Video S2**). Applying this strategy to NIPBL-WT and mutant proteins, we found that NIPBL-WT and mutants also exhibit two distinct low-mobility states (i.e., state 1 and state 2), as we have shown previously for different classes of chromatin-associated proteins (28) with one or two minor higher mobility states (states 3 and 4) for the mutants (**Figures 1F-G, S1C-F**). States 1 and 2 represent the motion of chromatin-bound proteins, while states 3 and 4 likely represent diffusing molecules that are transiently trapped within the focal volume (28). Binding in the lowest mobility state (state 1 in red - **Figures 1F-G, S1C-F**) requires an intact DNA-binding domain (DBD) as well as domains necessary to recruit coregulators and is related to the transcriptionally active state of TFs (28,29). Mutations in both C1 and/or C2 primarily reduced the fraction of state 1 NIPBL, which decreased from 47% for WT to 20% for C1^mut^, 15% for C2^mut^, and only 5% for C1+C2^mut^ (**Figures 1G, S1C-F**). We also found that NIPBL can transition between states 1 and 2 (Figure S1G) albeit with a preference to remain in the same state. Compared to WT, both C1^mut^ and C2^mut^ exhibit a lower probability to remain in state 1 and to transition from state 2 to state 1, whereas C1+C2^mut^ predominantly transitions into state 2 (**Figure S1G**). This suggests that interactions with co-regulators via C1 and C2 likely facilitate the transition of NIPBL into state 1.

Based on the above observations, we tested the effect of the LxxLL cluster mutations on the basal transcriptome. We performed RNA-seq in the cl15-derived cell lines (cl15-EV/WT/C1^mut^/C2^mut^/C1+C2^mut^, see Methods) upon endogenous NIPBL-KD. Cl15-EV cells expressing endogenous NIPBL served as a control. NIPBL-KD resulted in differential expression of 471 genes (**Figure 1H, S1H**), of which ∼74% (348) were rescued by ectopic NIPBL-WT (**Figure 1H**). The expression levels of the 348 NIPBL-regulated genes in cl15-WT were highly similar to those in cl15-EV cells expressing endogenous NIPBL, highlighting the quality of our rescue system (**Figure 1I**). In contrast, cl15-C1^mut^/C2^mut^/C1+C2^mut^ all had expression profiles like that of the NIPBL-KD cells, demonstrating that NIPBL LxxLL mutants are unable to rescue the transcriptional defects resulting from endogenous NIPBL-KD. These data show that C1 and C2 play an important role in the regulation of the basal transcriptome by NIPBL (**Figure 1I**). We note that even though C1^mut^ and C1+C2^mut^ were expressed 2-fold higher than WT (**Figure S1B**), they were unable to restore the basal gene expression program.

Taken together, these data show that C1 and C2 are important for NIPBL to associate with chromatin in general, but also for NIPBL to bind in or transition to state 1, which is associated with transcriptional activity. Notably, C1+C2^mut^ shows only marginal binding in state 1 (**Figure 1G**). Mutations in both C1 and C2 significantly alter the basal transcriptome, and we hypothesized that this might be because of the role of these Leu-rich clusters in mediating NIPBL-co-regulator interactions necessary to maintain gene expression.

### Chromatin-associated proteins interact with NIPBL-MAU2 in an LxxLL-dependent manner

To identify proteins that interact with NIPBL on chromatin in an LxxLL-dependent manner, we performed co-immunoprecipitation followed by mass spectrometry specifically on the chromatin-associated fraction of our cl15-WT, cl15-C1^mut^, and cl15-C2^mut^ cell lines (**Figure 2A**). NIPBL’s chromatin interactome included known partners such as MAU2, STAG2, and RAD21. Remarkably, the NIPBL interactors included TFs, chromatin remodelers, and cohesin, integrator and RNA Pol II subunits (**Figure 2A**). Several of the NIPBL interactors were lost in C1^mut^ and C2^mut^, suggesting that both clusters might help recruit an assortment of nuclear proteins (**Figure 2A**).

**Figure 2:**
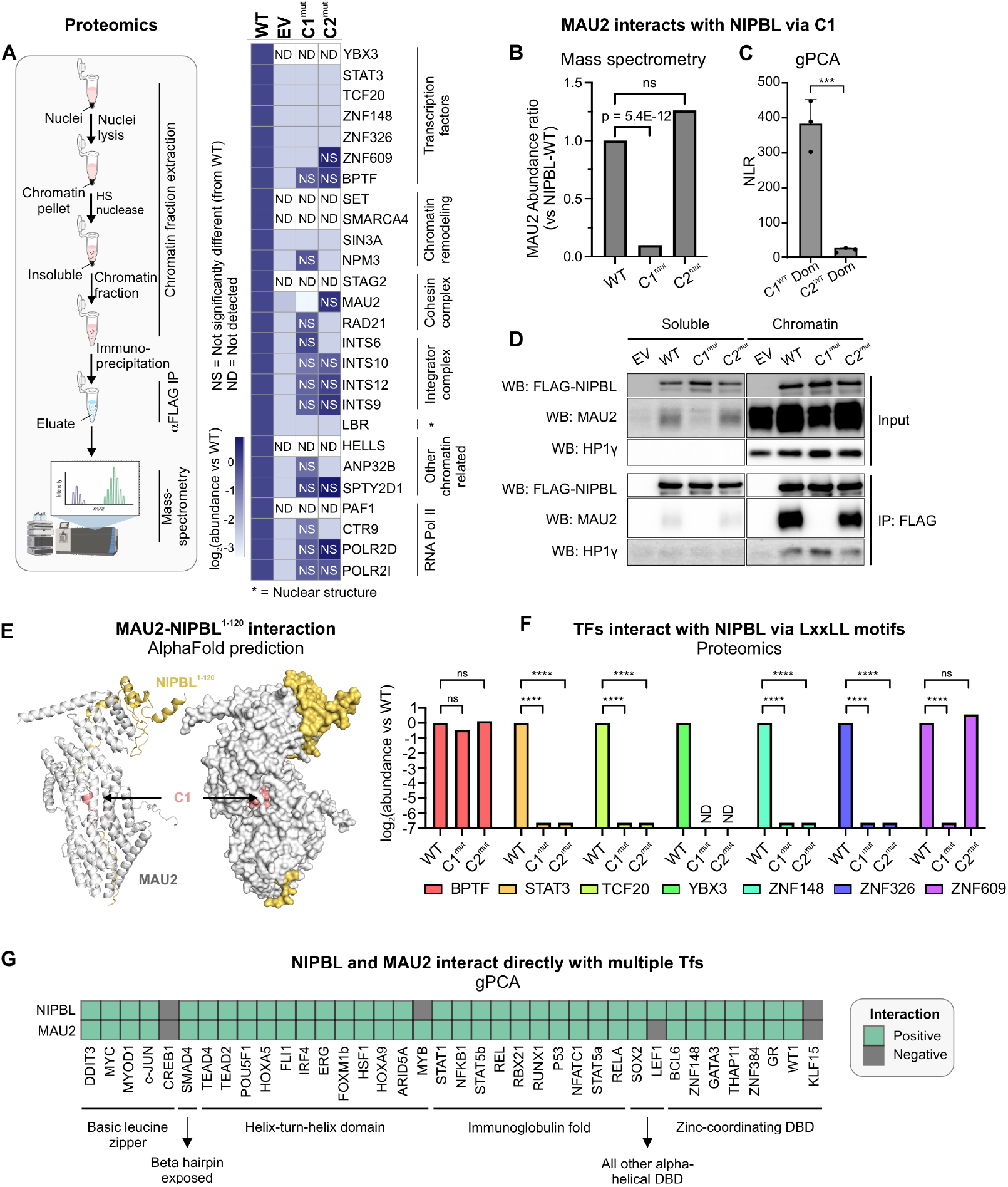
Leu-rich clusters recruit diverse chromatin-associated proteins. **(A)** (Left) Schematic representation of the proteomics experiments. (Right) Heatmap of abundance ratio of peptides detected in WT vs indicated NIPBL mutants. **(B)** MAU2 abundance detected in NIPBL-C1^mut^ and C2^mut^ relative to WT. p-value is reported from a one-way ANOVA. **(C)** NLRs for MAU2 against mNIPBL-C1^WT^ and mNIPBL-C2^WT^ domains. Error bars denote standard deviation. ^***^p<0.001 (paired t-test). **(D)** Co-immunoprecipitation of MAU2 with NIPBL-WT, C1^mut^, and C2^mut^ using the anti-FLAG antibody. **(E)** AlphaFold2-Multimer prediction of mNIPBL^1-120^ (yellow) with mMAU2 (silver). C1 is indicated in salmon. **(F)** Mass-spec log_2_(abundance ratio: C1^mut^/C2^mut^ vs WT) of selected TFs. ^****^p<0.001 (one-way ANOVA). **(G)** Heatmap of gPCA interactions of NIPBL and MAU2 against a panel of TFs. See also, Figure S2.

To validate our proteomics data, we next used the gaussia protein-fragment complementation assay (30) (gPCA, see Methods) (**Figure S2A**). As expected, full length NIPBL-WT scored positively (normalized luminescence ratio [NLR]>3.5) with MAU2, as well as with other cohesin subunits (SMC1A, SMC3, and RAD21). As a control, CTCF, which does not interact with NIPBL (31), scored negatively (**Figure S2A**). MAU2 was specifically lost only in C1^mut^ but not C2^mut^, suggesting that the NIPBL-MAU2 interaction could occur through C1 (Figures 2B). Indeed, the isolated C1^WT^ region of NIPBL (mouse aa13-120) was able to interact with MAU2, in contrast to the C2^WT^ region (mouse aa1485-1783) (**Figure 2C**). This is consistent with previous work that localized the MAU2 interaction interface of NIPBL to aa1-38 (human) (32). Chromatin-associated MAU2 co-immunoprecipitated with NIPBL-WT and C2^mut^ but not with C1^mut^ (**Figure 2D**), providing further evidence that while on chromatin, NIPBL interacts with MAU2 through C1. HP1-γ, which is known to interact with NIPBL through a PxVxL motif (33) (mouse aa995-999), was used as a positive control, and was unaffected by the C1 and C2 mutations (**Figure 2D**).

We next used the latest version of AlphaFold2-Multimer (34) to predict the structure of a complex consisting of full-length MAU2 and the first 120 residues of NIPBL (NIPBL^1-120^, see Methods). AlphaFold2-Multimer predictions showed a barrel-like complex where the disordered region of NIPBL is buried within MAU2. This arrangement minimizes solvent exposure, stabilizing NIPBL^1-120^, and reveals how C1 directly interacts with MAU2 (**Figure 2E**). Together, these data show that NIPBL directly interacts with MAU2 via its C1 region through the LxxLL motifs.

After obtaining the structure of the mouse MAU2-NIPBL complex, predicted by AlphaFold2-Multimer, we observed that its yeast homologs, Scc4-Scc2, adopt a remarkably similar conformation, as seen in the crystal structure (PDB: 4XDN) (35). In both structures, MAU2/Scc4 (gray) encases the N-terminal region of NIPBL/Scc2, stabilizing its extended conformation within the axial groove (**Figure S2B**). This suggests a conserved mechanism of interaction between species.

Remarkably, 7 TFs were found to interact with NIPBL on chromatin through C1, C2, or alternative surfaces (**Figures 2A, F**). We used gPCA to confirm that ZNF609 (previously shown to interact with NIPBL (16)), TCF20, and ZNF148 interact directly with NIPBL-WT (**Figure S2C**). These data lead to two non-exclusive hypotheses: (1) Since C1 is necessary for the MAU2-NIPBL complex formation and many TF-NIPBL interactions are lost in C1^mut^, MAU2 could bridge interactions between NIPBL and TFs. (2) The direct interaction between NIPBL and some TFs (e.g., TCF20, ZNF148, and ZNF609) and the loss of several TF-NIPBL interactions in C2^mut^ (**Figure 2A**) suggest that a set of TFs could directly interact with NIPBL through C2 in a MAU2-independent manner.

To examine the potential for direct TF-NIPBL and TF-MAU2 interactions, we tested a library of 38 representative TFs belonging to 6 TF superfamilies against NIPBL and MAU2 using gPCA. Strikingly, 34 TFs, spanning all the TF families, scored positively for interactions with both NIPBL and MAU2, including GR (which we have previously shown to interact with NIPBL (7)) (**Figures 2G, S2D-E**).

These data show that several TFs can interact with both NIPBL and MAU2 and suggest that TFs may form a ternary complex with the NIPBL-MAU2 heterodimer.

### SRs associate with C2 via their LBDs

Since our proteomics data show that several TF-NIPBL interactions are lost in C2^mut^ (**Figure 2F**), we ran a gPCA screen against an isolated C2 domain (mouse aa1485-1783) and found that 11 of the 38 tested TFs scored positively (**Figure 3A**), indicating that the C2 domain is sufficient to drive interactions between some TFs and NIPBL (see Limitations). To determine the interaction interfaces between NIPBL/MAU2 and TFs, we next focused on NRs for a more detailed characterization.

**Figure 3:**
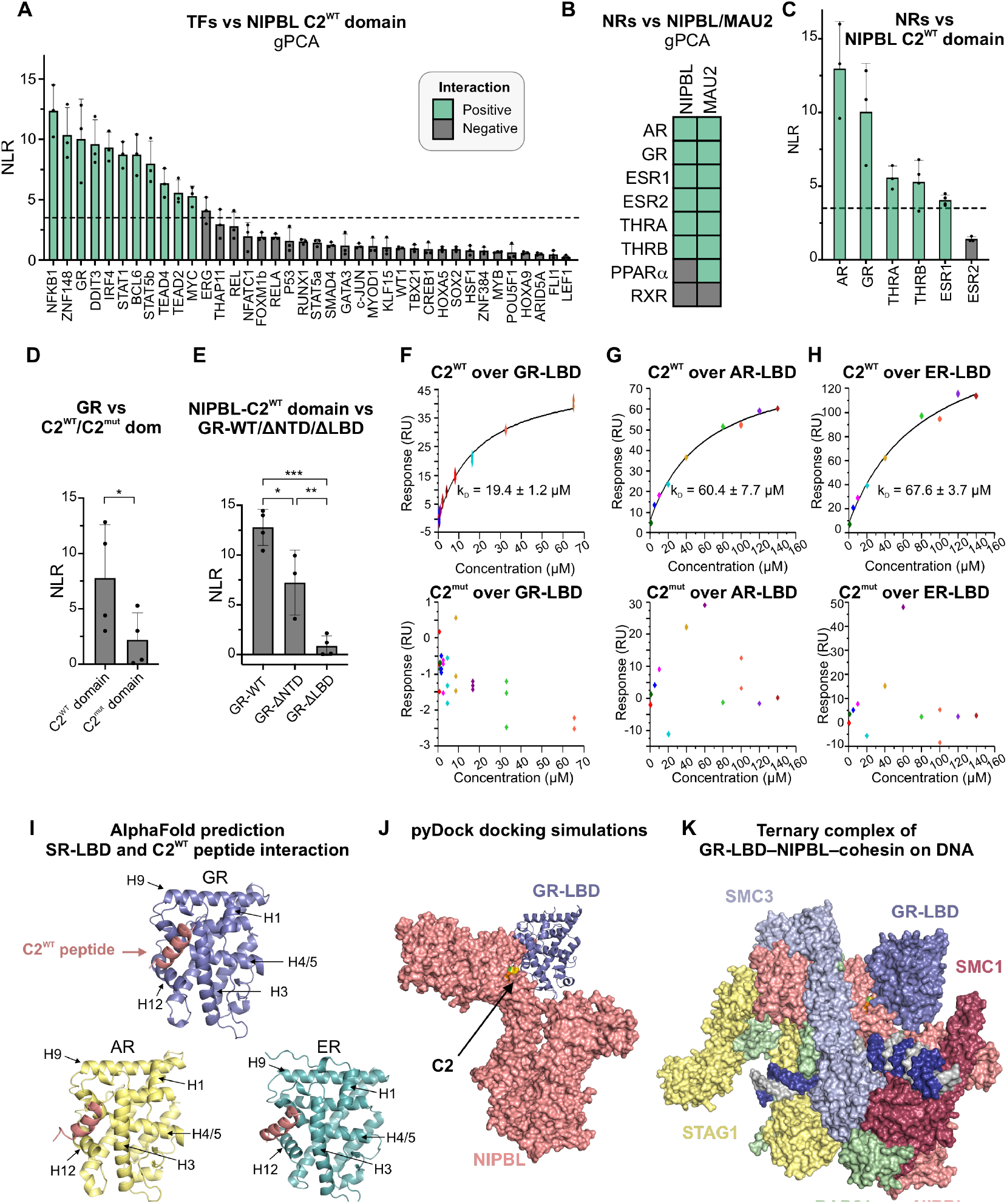
SR-LBDs interact with NIPBL through C2. **(A)** NLRs of the NIPBL-C2^WT^ domain against indicated TFs. Green = positive and black = negative interactions. **(B)** Heatmap of interactions between NIPBL/MAU2 and NRs detected by gPCA. **(C-E)** NLRs for **(C)** NIPBL-C2^WT^ domain against the NRs that scored positively against NIPBL-WT; **(D)** GR against NIPBL-C2^WT^/C2^mut^ domains; ^*^ p<0.05 (paired t-test); **(E)** NIPBL-C2^WT^ domain vs GR-WT, GR lacking its NTD (GR-ΔNTD), or its LBD (GR-ΔLBD), ^*^p<0.05, ^**^p<0.01,^***^p<0.001 (one-way ANOVA and Tukey test for multiple comparisons). Error bars in panels A-E represent the standard deviation across measurements. **(F-H)** SPR sensorgrams for indicated human (h) SR-LBDs against human NIPBL-C2^WT^ (top) and C2^mut^ (bottom) peptides. **(I)** AlphaFold2-Multimer predictions for the interaction between hNIPBL-C2^WT^ peptide (salmon) and the LBDs of hGR, hAR, and hER. **(J)** Docking prediction for the hGR-LBD (blue) with the structured portion of hNIPBL (salmon). **(K)** Superposition of the pyDock prediction of the NIPBL-GR-LBD structure with the cryo-EM structure of the NIPBL-cohesin-DNA complex (24). See also, Figure S3.

Using gPCA, we found that 7 out of 8 tested NRs interact with MAU2 and 6 with NIPBL-WT (**Figures 3B, S3A-B**). Notably, 5 of these 6 NRs also scored positively when tested against the NIPBL-C2 domain, including all but one SR (**Figure 3C**). Focusing on GR, AR, and ER as representative SRs, we found that compared to C2^WT^, their interactions with the C2^mut^ domain were largely abolished, strongly suggesting that GR, AR, and ER interact with the C2 region of NIPBL through the LxxLL motifs (**Figures 3D, S3C-D**). SRs consist of a disordered NTD, and structured DBD and LBD. To locate the C2-interacting surface within GR, we generated GR constructs lacking either the LBD (GR-ΔLBD) or the NTD (GR-ΔNTD). While GR-ΔNTD interacts with C2^WT^, GR-ΔLBD does not, showing that a region spanning the GR-LBD and DBD is necessary and sufficient to mediate the interaction with the NIPBL-C2 domain (**Figure 3E**).

We next used SPR to measure the binding affinity of the GR, AR, and ER LBDs with short peptides spanning the C2^WT^ or C2^mut^ cluster (**Figure S3E, see Methods**). The C2^WT^ peptide has a helical conformation, like that of the C2 cluster within NIPBL (**Figure S3F**). All three SR-LBDs showed positive interactions with the C2^WT^ peptide with micromolar affinities (**Figure 3F-H, top**), while no interactions were detected with the C2^mut^ peptide (**Figure 3F-H, bottom**), demonstrating that SR-LBDs can engage in specific interactions with the LxxLL motifs within the C2 region of NIPBL.

High-confidence models from AlphaFold2-Multimer for all three SRs predicted that the NIPBL-C2^WT^ peptide aligns closely with conserved hydrophobic residues of the AF-2 pocket of the SR-LBD (**Figures 3I, S3G**). The AF-2 pocket is a hydrophobic groove formed by helices H3, H5, and H12 upon ligand binding and its role in recruiting co-regulators that modulate SR transcriptional activity has been extensively characterized (36-38).

Based on these data, we modeled the interaction between the GR-LBD and the structured portion of NIPBL using pyDock (39). The resulting docking shows how the AF-2 surface of the GR-LBD has the potential to interact with NIPBL through the C2 LxxLL motif cluster (**Figure 3J**). Next, we superimposed the GR-LBD-NIPBL docking prediction (**Figure 3J**) on the recent cryo-EM structure of the NIPBL-cohesin-DNA complex (24) (**Figure 1D**) to check whether they are mutually compatible. The GR-LBD can bind to the C2 LxxLL motif cluster without steric hindrance from any of the other cohesin complex subunits, while the orientation of the GR-LBD in this structure allows sufficient space for the GR-DBD to bind to DNA (**Figure 3K**).

We next sought to investigate the GR-MAU2 interaction surfaces and examine the consequences of the C1 and C2 mutations on GR-mediated gene expression.

### C1 and C2 are necessary for the formation of a NIPBL-MAU2-GR ternary complex and to maintain the GR-regulated transcriptome

First, we observed that the GR-MAU2 interaction also depends on a region spanning the GR-DBD and LBD (**Figure 4A**). We then used AlphaFold2-Multimer to predict possible conformations of the NIPBL^1-120^-MAU2-GR-LBD complex. We found that the GR-LBD interacts with MAU2 through helix 9, leaving the AF-2 pocket available for interactions with NIPBL-C2, while preserving the MAU2-C1 assembly (**Figures 2E, 4B**). NIPBL, MAU2, and GR co-immunoprecipitated in cl15-WT cells (**Figure 4C**), strongly supporting the existence of a GR-NIPBL-MAU2 ternary complex *in vivo*.

**Figure 4:**
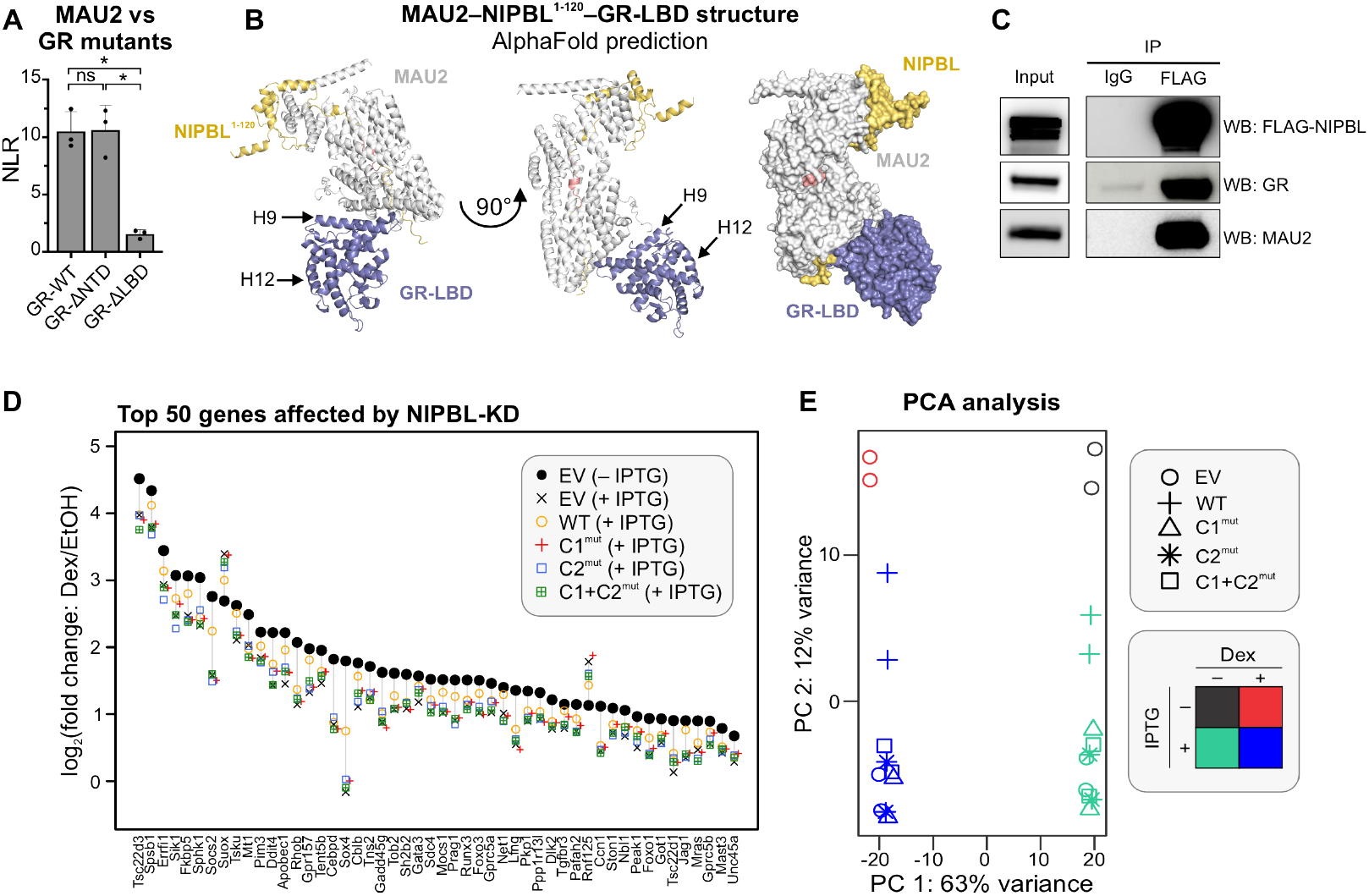
GR forms a ternary complex with NIPBL/ MAU2 with implications for glucocorticoid signaling. **(A)** gPCA results of MAU2 against GR-WT, GR-ΔNTD, and GR-ΔLBD. ^*^ p<0.05 (one-way ANOVA followed by Tukey test). Error bars denote the standard deviation. **(B)** AlphaFold2-Multimer prediction of the composite structure of mMAU2-mNIPBL^1-120^-mGR-LBD. **(C)** Triple-IP of FLAG-NIPBL, MAU2, and GR in cl15-WT cells. **(D)** Scatter plot of the top 50 genes whose Dex-response is affected upon NIPBL-KD. **(E)** PCA plot of the RNA-seq data across multiple conditions presented in Figures 1 and 4. See also, Figure S4.

To test if the formation of the GR-NIPBL-MAU2 ternary complex plays a role in GR-mediated transcription, we profiled transcriptional changes in cl15-EV/WT/C1^mut^/C2^mut^/C1+C2^mut^ cells after dexamethasone (Dex) treatment. Since not all GR-regulated genes are NIPBL-dependent, we focused on the top 50 genes whose Dex-induction was affected by NIPBL-KD. Across all these genes, the Dex response of cl15-WT cells was the closest to that of control EV cells expressing endogenous NIPBL. Strikingly, none of the C1/C2 mutants performed as well as NIPBL-WT. For most genes, the transcriptional response was similar to that of the cl15-EV control cells with endogenous NIPBL-KD, indicating their inability to compensate for the loss of endogenous NIPBL (**Figures 4D-E, S4**). These data provide a window into the functional consequences of NIPBL-TF interactions through the lens of glucocorticoid signaling: Interaction with NIPBL is important to maintain robust gene activation by TFs, at least in response to GR.

Altogether, our biochemistry, imaging, and structural prediction data show that GR (and most probably other TFs), MAU2, and NIPBL can form a ternary complex *in vivo* to regulate gene expression.

## DISCUSSION

Long-range chromatin interactions are the bedrock of genome organization and the NIPBL-cohesin complex is instrumental in the formation of dynamic enhancer-promoter loops. While NIPBL is commonly believed to localize cohesin at enhancers (7,15,31), it lacks the ability to identify specific DNA sequences. In our previous work, we had proposed that GR and other TFs can interact with NIPBL to localize cohesin to their target enhancers (7). Here, we identified two clusters of Leu-rich motifs within NIPBL (C1 and C2) whose mutations affect NIPBL dynamics and the basal transcriptome (**Figure 1**). Our proteomics data show that C1 and C2 serve as platforms for the recruitment of diverse chromatin-associated proteins, notably TFs (**Figures 2A, B, F**). Using orthogonal methods, we showed that the C1 motif is necessary for the MAU2-NIPBL interaction (**Figure 2B-E**) and that MAU2 adopts a barrel-like configuration engulfing C1 in its axial groove (**Figures 2E, 5A**). Remarkably, 34 out of 38 tested TFs can directly interact with NIPBL and MAU2 (likely through different mechanisms), laying the foundation for the proposed ternary complex model (**Figures 2G**). Focusing on SRs, we showed that GR, AR, and ER interact with the C2 region of NIPBL through their LBDs and in an LxxLL motif-dependent manner (**Figures 3**). Narrowing down to GR, we then predicted that GR interacts with NIPBL-C2 through its AF-2 pocket (**Figures 3J, 5A**), and with MAU2 via helix 9 in its LBD (**Figures 4B, 5A**). Mutations in C1 and/or C2 result in an impaired Dex-response (**Figures 4D-E, S4**), underscoring the importance of these Leu-rich motifs for the regulation of a subset of GR-responsive genes.

The GR-LBD crystallizes as tetramers (40) and higher oligomers (41) *in vitro. In vivo*, full-length GR is a dimer in the nucleoplasm and a tetramer on chromatin (42). The docking and AlphaFold2-Multimer predictions demonstrate pair-wise interactions between the GR-LBD and C2, and the GR-LBD and the MAU2-NIPBL^1-120^ complex (**Figure 5A**). Both the NIPBL-C2/GR-LBD interaction through the AF-2 pocket, and the MAU2/GR-LBD interaction through helix 9 do not interfere with the canonical GR dimerization interfaces that we have identified previously (40). As such, at least three distinct possibilities exist for the configuration of the GR-NIPBL-MAU2 ternary complex *in vivo*: (1) A single GR molecule interacts with both NIPBL (via the AF-2 pocket) and MAU2 (via helix 9) (**Figure 5B, left**), (2) a GR dimer interacts with the NIPBL/MAU2 complex through a single GR molecule (**Figure 5B, middle**), or (3) within a GR dimer, one GR molecule interacts with NIPBL (via the AF-2 pocket) and the second GR molecule interacts with MAU2 (through helix 9) (**Figure 5B, right**). GR, thus, has multiple geometrically compatible configurations to form a ternary complex with the NIPBL/MAU2 heterodimer.

**Figure 5:**
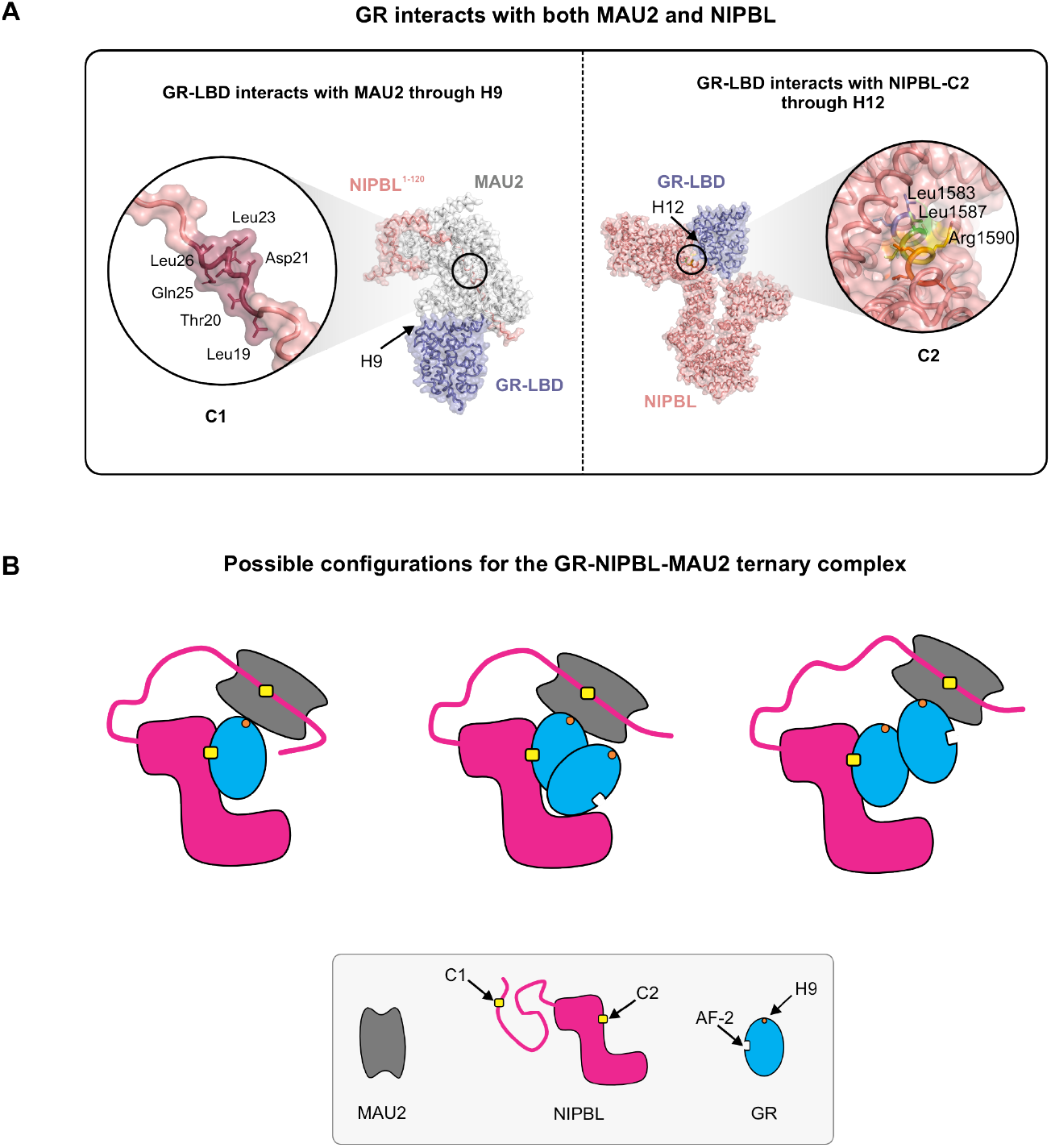
Possible models for a GR-NIPBL-MAU2 ternary complex. **(A)** (Left) The glucocorticoid receptor ligand-binding domain (GR-LBD) interacts with the NIPBL^1-120^-MAU2 complex through helix 9. NIPBL^1-120^ and MAU2 interact through the C1 LxxLL motif cluster. (Right) The GR-LBD interacts with NIPBL-C2 through helix 12 in the AF-2 pocket. Key residues within C1 and C2 are indicated in the insets. **(B)** Possible configurations of the GR-NIPBL-MAU2 ternary complex: (Left) A single GR monomer can interact with both NIPBL-C2 through the AF-2 pocket and MAU2 through helix 9. (Middle and Right) Since the dimerization interface of GR is not occluded by either NIPBL or MAU2, a GR dimer can interact with NIPBL/MAU2 either through (Middle) a single GR molecule or (Right) with one GR molecule interacting with NIPBL-C2 and another one interacting with MAU2.

NIPBL is an essential component of the loop extrusion machinery and mutations in NIPBL are responsible for almost 70% of Cornelia de Lange syndrome (CdLS) cases (43). CdLS is a rare genetic disorder characterized by developmental defects, facial abnormalities, gastroesophageal, heart and ocular dysfunction, and cognitive impairments (43,44). Understanding the molecular mechanism behind this disease is of fundamental importance to devise therapeutics. Strikingly, CdLS mutations have been localized to specific Leu residues in C1 (p.Leu23_Gln25del)(45) and C2 (p.Leu1584Arg)(46) and could impair cell growth as has been shown in yeast (20). Since C1 and C2 serve as platforms for the direct (via C2) or indirect (via C1-MAU2) recruitment of multiple TFs, our data suggest that these mutations could result in the loss of co-regulators and a possible mislocalization of NIPBL on chromatin, and hence, alter cohesin binding.

In this study, we proposed a ternary complex formed by GR with the NIPBL-MAU2 heterodimer and investigated the structural underpinnings of this assembly. Moreover, our SPR and gPCA data for AR and ER suggest that this model may extend to other SRs and maybe even other NRs (such as THRA and THRB). However, it would be naïve to expect a universal mechanism for all TFs. Even though our gPCA data strongly indicate that many families of TFs can interact with both NIPBL and MAU2, the precise mechanisms are likely to differ. Unraveling the mechanisms of all these interactions will be the subject of our future work.

## LIMITATIONS

The challenges involved in detecting and knocking down NIPBL have been an open secret in the field (13). To circumvent the heterogeneity in siRNA-based knockdown, we generated a clonal cell line expressing an IPTG-inducible shRNA against NIPBL. Despite harboring an inducible shRNA, after 72 h of IPTG treatment, we detected ∼70% depletion of *Nipbl* mRNA (**Figure S1A**). While we acknowledge that this knockdown isn’t perfect, this reagent represents our best attempt at developing a NIPBL-KD cell line in the 3134 mouse mammary adenocarcinoma cell line. The subtle yet reproducible effects of NIPBL-KD on the basal and GR-mediated transcription are likely consequences of incomplete NIPBL-KD.

Since high-throughput detection methods of PPIs can only capture ∼20-30% of PPIs (47), the proteins detected in our mass-spec experiment likely represent a small fraction of proteins that indeed interact with NIPBL *in vivo*. We also recognize that PPIs are highly dynamic and do not result in static structures. The TF-MAU2 interaction and TF-NIPBL interaction need not occur simultaneously.

gPCA is a cell-based *in vitro* assay that identifies pairs of proteins that have the potential to interact in cells. We validated this screen using multiple controls. A positive interaction can be taken at face value. However, a negative interaction must be validated by testing all 4 configurations of the split luciferase (N- and C-terminus) on each protein.

## ACKNOWLEDGMENTS

We would like to thank the Viral Vector Platform of the GIGA, ULiège for the production of lentiviral particles used for the NIPBL-KD, the GIGA Flow Cytometry Core for cell sorting, Luke Lavis (Janelia Research Campus) for the Halo-JF dyes, Razi Raziuddin and Le Hoang (NCI) for assistance with experiments, and Supriya Vartak for constructive feedback.

## AUTHOR CONTRIBUTIONS

Conceptualization, G.F.; Methodology, G.F., K.W., D.A.S., A.J.-P., E.E.-P., F.D., and J.F.-R.; Software, K.W., A.J.-P., S.K., D.A.B., and J.F.-R.; Formal Analysis, G.F., K.W., A.J.-P., S.K., D.V., and A.U.; Investigation, G.F., K.W., D.A.S., A.J.-P., M.L., A.A.-M., T.A.J., M.K., L.W., and L.R.; Writing – original draft, K.W.; Writing – review and editing, G.F., K.W., D.A.S., A.J.-P., A.A.-M., E.E.-P., F.D., and G.L.H; Visualization, G.F., K.W., D.A.S., A.J.-P., S.K., and A.A.-M.; Supervision, T.S.K., E.E.-P., J.F.-R., F.D., and G.L.H.; Project administration, F.D., and G.L.H; Funding acquisition, T.S.K., A.U., E.E.-P., J.F.-R., F.D., G.L.H.

## DATA AVAILABILITY

## Materials availability

All plasmids and cell lines generated in this study will be available upon request to the corresponding authors, Gordon L. Hager and Franck Dequiedt.

## Data and code availability

- SMT data are available in Zenodo (<u>doi.org/10.5281/zenodo.14277999</u>)
- SMT software is available at Zenodo (<u>doi.org/10.5281/zenodo.7558712</u>)
- pEMv2 software is available at https://github.com/MochrieLab/pEMv2

## MATERIALS AND METHODS

### Cell lines

HEK293T cells and cell lines derived from the 3134 mouse mammary adenocarcinoma cell line were grown in Dulbecco’s modified Eagle’s medium (DMEM) and 10% fetal bovine serum (FBS) or char-coal-stripped FBS supplemented with sodium pyruvate, L-glutamine, and non-essential amino acids.

### Plasmids and cloning

shRNA lentiviral plasmids were purchased from VectorBuilder. These plasmids allow IPTG-inducible shRNA expression for targeting mouse *Nipbl* (GATTGTGGAGAGACCTAATTA, VB220804-1048fvd) driven by a U6/2xLacO promoter along with *LacI* and puromycin resistance expression under the control of mPGK promoter. Vectors containing 3xFLAG-Halo mouse NIPBL WT, C1^mut^, C2^mut^, and C1+C2^mut^ were custom-made by Thermo Fisher with an shNIPBL insensitive sequence, then cloned into a PiggyBac carrier plasmid using NheI and NotI restriction sites.

Most ORFs used for the gPCA experiments were already inserted in an entry vector and directly retrieved from the hORFeom7.1 and 8.1 collections (48) (Tables S1-S2). Other ORFs were inserted into a pDONR223 by PCR amplification followed by BP reaction (Gateway recombination technology, Invitrogen) (see Table S1,S2 for details). All ORFs clones were sequenced. pDONRs containing ORFs were then cloned in destination vectors containing GLucN1 and GLucN2 fragments of the *Gaussia princeps* luciferase (49) by LR Clonase reaction.

### Lentiviral particle production

Lentiviral vectors were generated by the GIGA Viral Vectors platform (University of Liège, Belgium). Briefly, Lenti-X 293T cells (Clontech®, 632180) were co-transfected with a pSPAX2 (Addgene®, Cambridge, MA, USA) and a VSV-G encoding vector (50). Viral supernatants were collected at 96 h post transfection, filtered (0.2 *μ*m) and concentrated 100x using Lentivirus concentration solution (SanBio, TR30026). The lentiviral vectors were then titrated with qPCR Lentivirus Titration (Titer) Kit (ABM^®^, LV900, Richmond, BC, Canada). The absence of mycoplasma was confirmed using MycoAlert™ PLUS Mycoplasma Detection Kit (Lonza, LT07-710).

### Generation of a NIPBL knock-down and rescue system

3134 cells were transduced (MOI 800) with protamine sulfate. After 72 h, the cells were treated with 1 *μ*g/mL puromycin for another 72 h and then amplified. A single-cell clone (clone 15 [cl15]) where *Nipbl*-KD was above 70% by qPCR after 72 h of 1 mM IPTG treatment (**Figure S1A**) was selected. Cl15 cells were then transfected with both a transposase and a PiggyBac carrier plasmid expressing the 3xFLAG-Halo conjugated NIPBL construct. A control cell line expressing only the 3xFLAG-Halo was also generated (referred to as the empty vector [EV] cell line). After selection with blasticidin (20 *μ*g/ml) for 72 h, the HaloTag was used to FACS sort the cells to obtain similar expression of the ectopic NIPBL variants.

### SMT sample preparation

For SMT experiments, 3134 cl-15 derived cl15-WT, cl15-C1^mut^, cl15-C2^mut^, and cl15-C1+C2^mut^ cells were plated in 2 well chamber slides (Cellvis, Mountain View, CA, USA) 24 hours prior to imaging. On the day of imaging, cells were incubated in phenol red-free complete medium containing 10-25 pM JFX650 dye (to visualize single molecules) and 100 pM JF549 (over-stained to mark the nucleus) for 20 min followed by 3 washes. The cells were washed once more after 10 min. The sample was mounted on the microscope and allowed to equilibrate for 20 min before data acquisition.

### Microscopy

All the SMT experiments were performed on a custom-built HILO microscope at the Optical Microscopy Core of the NCI (previously described in ref (51)). Briefly, the HILO microscope has a 150X 1.45 NA objective (Olympus Scientific Solutions, Waltham, MA, USA), a 561 nm laser (iFLEX-Mustang, Excelitas Technologies Corp., Waltham, MA, USA), a 647 nm laser (OBIS 647 LX, Coherent, Inc., Santa Clara, CA), and an acousto-optical tunable filter (AOTFnC-400.650, AA Optoelectronic, Orsay, France). An Okolab stage-top incubator maintains the temperature at 37°C and CO_2_ at 5% (Okolab, Pozzuoli NA, Italy). An EM-CCD camera (Evolve 512, Photometrics, Tucson, AZ, USA) with the gain set to 300 was used to collect images. The cell nucleus was centered and focused within the illumination field using the 549 channel. For the fast SMT protocol, images were collected in the 647 channel continuously every 12 ms for a total of 1500 frames. For the intermediate SMT protocol, images were collected every 200 ms with an exposure time of 10 ms for a total of 2 min (600 frames). The laser power was set at 0.96 mW. The pixel size for this setup is 104 nm.

For the fast SMT dataset (12 ms intervals), 70-142 cells were imaged per condition while for the intermediate SMT dataset (200 ms intervals), 53-67 cells were imaged per condition with at least two independent biological replicates each.

## SMT analysis

### Tracking

Single molecules were tracked using TrackRecord, described previously (52) and available at https://doi.org/10.5281/zenodo.7558712. The TIF-stacks were filtered using a band-pass filter. The sum projection of the time series was used to define a hand-drawn region of interest. A particle intensity threshold at which 95% of the detected particles had a signal-to-noise ratio of at least 1.5 was selected for the particle detection step. Raw positions of the particles were identified from the filtered image. These positions were fit to a two-dimensional Gaussian point spread function (PSF) for sub-pixel localization. The PSF fitting step was performed on the unfiltered image. Detected particles were connected to form tracks based on a nearest-neighbor algorithm with the following parameters: maximum jump = 4 pixels (416 nm), shortest track = 2 frames, gaps to close = 1 frame. These tracking parameters have been extensively validated in our previous work (28).

### Perturbation expectation maximization (pEMv2)

All tracks obtained from the 12 ms and 200 ms interval movies were analyzed using pEMv2(53) to identify the optimal number of mobility states that best describes the ensemble of trajectories. Tracks were split into sub-tracks of 7 frames as has been done previously (28,54). pEMv2 was allowed to explore between 1 and 15 states with 3 covariance matrix features. 50 reinitializations and 200 perturbations were performed for each step. The maximum number of iterations was 10,000 and the convergence criterion for the log-likelihood function was set to 10^−7^. Each sub-track was assigned to the state for which it had the highest posterior probability.

### Calculation of the bound fraction

Following the pEMv2 analysis, states with linear mean-squared displacements over lag-times were identified as the diffusive (unbound) states, while the other states were classified as bound. The number of bound particles is thus the number of sub-tracks detected in the ‘bound’ pEMv2 states multiplied by the length of each sub-track (which is 7 frames):

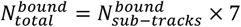

The bound fraction is thus calculated as:

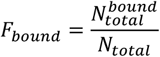

where *N*_*total*_ is the total number of detected particles.

### Calculation of pEMv2 state fractions and confidence intervals from the 200 ms data

To calculate the population fractions and representative MSD curves (**Figure S1C-F**), only sub-tracks for which the difference between the two highest posterior probabilities (ΔPP) was at least 0.2 (as has been done previously (28)) were considered. This was done to ensure that sub-tracks with similar posterior probabilities for more than one state were not included in the analysis. Further, states with a population fraction less than 5% were not retained in downstream analysis (**Figures S1C-F**). The population fraction for each remaining state was calculated as the fraction of subtracks with ΔPP ≥ 0.2 that belong to the respective state.

The confidence intervals of the population fractions were calculated using a bootstrapped analysis. 1000 bootstrapped ensembles of tracks were generated by resampling the tracks with replacement. The population fractions of each of the ensembles was calculated using the procedure described above. From this distribution of population fractions, the 95% confidence interval was calculated. The population fractions and confidence intervals were normalized to the bound fraction.

### Calculation of transition probabilities

Transition probabilities and associated statistical significance were calculated as described previously (28). All sub-tracks assigned to states other than states 1 and 2 were re-assigned to a third state named ‘Other’. For each track, the number of transitions between each pair of these three states was calculated. Only tracks containing 3 or more sub-tracks were analyzed. These counts were summed up to generate a transition matrix *T* where each element *T*(*i, j*) represents the number of transitions between states *i* and *j*. The transition matrix was normalized to calculate the transition probability matrix *P*_*t*_ where 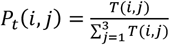.

To statistically test whether the calculated transition probabilities were different from those that would arise from a random ensemble with the same population fractions, 1000 randomized ensembles were generated by shuffling the sub-track labels. The fraction of these 1000 randomized trials with a transition probability 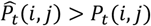 is reported as the statistical significance value.

### Immuno-precipitation on the chromatin fraction

3134 cells expressing the different variants of mNIPBL were plated 72 hours before the experiment, then trypsinized, washed with PBS, and the nuclei purified. Cells were then resuspended at 3 × 10^6^ cells/mL and incubated on ice for 10 min in Buffer A (Tris-Cl pH 8.0 15 mM, NaCl 15 mM, KCl 60 mM final, EDTA 1 mM final, EGTA 0.5 mM final) and nuclei were pelleted by centrifugation at 350 g for 5 min. The nuclei were then washed with the NSB buffer (Glycerol 25%, MagAc2 5 mM, HEPES pH 7.3 5 mM, EDTA 0.08 mM, DTT 0.5 mM) and lysed in lysis buffer (glycerol 10%, HEPES pH 7.5 20 mM, NaCl 150 mM, MgCl2 1.5 mM, NP40 0.8%, DTT 0.5 mM) by 30 min of rotation at 4°C. Lysates were further centrifuged for 5 min at 9000 g and the supernatants collected as the nucleoplasmic fractions. The pellets were then resuspended in the lysis buffer complemented with 100 units/mL of HS-Nuclease (MoBiTec) and incubated at room temperature (RT) for 1 h with rotation. Resulting nuclear lysates were then centrifuged for 5 min at 9000 g and the supernatants collected as the chromatin fractions. These fractions were further incubated with pre-washed antibody-charged magnetic beads overnight at 4°C. Beads were then washed twice with the lysis buffer and three times with the wash buffer (glycerol 10%, HEPES pH7.5 20 mM, NaCl 250 mM, MgCl2 1.5 mM, NP40 0.8%, DTT 0.5 mM). The magnetic beads-bound proteins were eluted with 80 *μ*g/mL of 3xFLAG peptide (ApexBio) diluted in TBS for 1 h at 18°C and shaken at 1000 rpm. Eluates were then subjected to the western blot procedure or sent for proteomic analysis (**Figure 2A**). All buffers were supplemented with 1x protease inhibitor (Roche).

### Mass spectrometry

Protein eluates were loaded on a 10% acrylamide SDS-PAGE gel and subjected to a short migration of 15 min. The single protein band was visualized by colloidal Coomassie Blue staining and in-gel digested with Trypsin (Promega). Peptides were extracted with 0.1% trifluoroacetic acid in 65% acetonitrile and dried in a speedvac. Peptides were dissolved in solvent A (0.1% trifluoroacetic acid in 2% acetonitrile), directly loaded onto reversed-phase pre-columns (Acclaim PepMap 100, Thermo Fisher Scientific) and eluted in backflush mode. Peptide separation was performed using a reversed-phase analytical column (Easy-spray PepMap RSLC, 0.075 × 250 mm, Thermo Fisher Scientific) with a linear gradient of 4%-27.5% solvent B (0.1% formic acid in 80% acetonitrile) for 40 min, 27.5%-50% solvent B for 20 min, 50%-100% solvent B for 3 min, and holding at 100% for the last 10 min at a constant flow rate of 300 nL/min on an Ultimate 3000 RSLC system.

The peptides were analyzed by an Orbitrap Exploris240 mass spectrometer (Thermo Fisher Scientific). The peptides were subjected to NSI source followed by tandem mass spectrometry (MS/MS) coupled online to the nano-LC. Intact peptides were detected in the Orbitrap at a resolution of 60,000. Peptides were selected for MS/MS using HCD setting at 30, ion fragments were detected in the Orbitrap at a resolution of 15,000. A data-dependent procedure that alternated between one MS scan followed by 40 MS/MS scans for ions above a threshold ion count of 1.0×10^4^ in the MS survey scan with 30.0 s dynamic exclusion. MS1 spectra were obtained with an automatic gain control target of 4×10^5^ ions and a maximum injection time set to auto, and MS2 spectra were acquired with an automatic gain control target of 5×10^4^ ions and a maximum injection set to auto. For MS scans, the m/z scan range was 350 to 1800.

The resulting MS/MS data were processed using the Sequest HT search engine within Proteome Discoverer 2.5 SP1 against a mouse protein database obtained from UniProt (55,520 entries, January 2023). Trypsin was specified as the cleavage enzyme allowing up to 2 missed cleavages, 4 modifications per peptide. Mass error was set to 10 ppm for precursor ions and 0.03 Da for fragment ions. Oxidation on Met (+15.995 Da), conversion of Gln (−17.027 Da) or Glu (−18.011 Da) to pyro-Glu at the peptide N-term, acrylamide modification of Cys (+71.037 Da) and phosphorylation on Ser, Thr or Tyr (+79.966 Da) were considered as variable modifications. False discovery rate (FDR) was assessed using Percolator and thresholds for protein, peptide, and modification sites were specified at 1%. Label-free quantification was performed within Proteome Discoverer using the area under the curve, normalization was performed based on the total amount of the 3xFLAG-HaloTag sequence contained in each sample.

The heatmap displaying chromatin-related proteins from the IP-MS proteomic experiment was generated using a custom R script. Abundance ratios were calculated relative to the NIPBL-WT condition. Proteins with abundance ratios not significantly different from NIPBL-WT (P-value ≥ 0.05) are labeled as ‘NS’ (not significant). Proteins that were undetected in a specific condition but detected in the WT samples are marked as ‘ND’ (not detected).

### Gaussia protein-fragment complementation assay (gPCA)

HEK293T cells were sub-cultured in 24-well plates and transfected with the appropriate constructs. After 48 h, cells were washed with PBS and processed according to the manufacturer’s instructions (Renilla Luciferase Kit, Promega). Lysates were then used to quantify luminescence in triplicate on a TriStar^2^ S LB 942 luminometer (Berthold). The remaining volume of lysate was used to assess the expression of the constructs by western blot analysis. At least three independent biological replicates were performed for every reported construct.

For each replicate, we calculated the Normalized Luminescence Ratio (NLR).

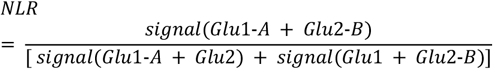

With A and B being the proteins of interest and the Glu1 and Glu2 the split luciferase fragments without any conjugated protein and Glu1-A and Glu2-B are Glu1 and Glu2 conjugated to proteins A and B respectively. Assessment of the interaction between two ORFs was done by performing a one-tail one-sample t-test against a value of 3.5 as suggested by Cassonnet et al (49). Positive interactions were identified with a p-value threshold of 0.05. Pairwise comparisons were performed when all the constructs were expressed to similar levels. Statistical significance was assessed either by a paired t-test or a one-way ANOVA followed by a Tukey test, as indicated in the legend.

### Peptides and proteins for surface plasmon resonance

Two peptides corresponding to residues 1576-1585 (cluster C2) of the human NIPBL protein were custom synthesized at Pepmic:

C2^WT^ (KPEWPAAELLLSLLGRLLVHQFSNKSTE), C2^mut^ (KPEWPAAEAAASAAGRAAVHQFSNKSTE).

Recombinant ancient GR-LBD (ancGR2-LBD, corresponding to residues 529-777 of the human receptor) cloned into a pMALCH10T vector was expressed as a fusion protein with an N-terminal maltose-binding protein (MBP) and a hexahistidine tag, and purified to homogeneity using standard chromatographic procedures. Recombinant AR-LBD (residues 672-920 of the human receptor), and ERα-LBD (residues 302-595 of the human receptor) cloned into pGEX vectors were expressed and purified following the same protocols as that for the ancGR2-LBD.

### Surface plasmon resonance

SPR analyses were performed at 25ºC in a BIAcore T200 instrument (GE Healthcare). Highly purified, dexamethasone-bound recombinant ancGR2-LBD, di-hydrotestosterone-bound AR-LBD, and estradiol-bound ER-LBD were diluted in 10 mM sodium acetate, pH 5.0, and directly immobilized on CM5 chips (GE Healthcare) by amine coupling at densities between 100 and 200 resonance units (RU). As a reference, one of the channels was also amine-activated and blocked in the absence of protein. Increasing concentrations of C2^WT^ and C2^mut^ from 0 to 160 *μ*M were run over the immobilized proteins. The running buffer was PBS, 1x Tween-20, 5% DMSO. Sensorgrams were analyzed with the BIAcore T200 Evaluation software 3.0 and fitted according to the Langmuir 1:1 model.

### AlphaFold predictions and analysis

Protein-protein complex predictions were performed using AlphaFold2-Multimer (34) to model the interactions of (1) the human NIPBL C2^WT^ helix with human SR-LBDs, (2) mNIPBL C2^WT^ domain with mMAU2, and (3) the ternary complex consisting of mGR-LBD, mNIPBL C1^WT^ domain, and mMAU2. For each of these three complexes, we executed five independent trajectories with each of the three available versions of AlphaFold2-Multimer (Version 1: v2.1.0, Version 2: v2.2.0, Version 3: v2.3.0), ensuring robust sampling of the conformational space. All generated structures were subjected to additional minimization steps using OpenMM to refine the models and improve energy stabilization. This resulted in a total of 330 models per complex (5 trajectories × 3 AlphaFold2-Multimer versions × 22 models per trajectory).

To select the best possible predictions, models were ranked based on a composite scoring function derived from AlphaFold’s internal model confidence metrics. Specifically, we used a weighted sum of the model’s per-residue confidence (ipTM score) and the overall structure confidence (pTM score), with the formula: *model confidence* = 0.8 × *ipTM* + 0.2 × *pTM*. Only models with a combined confidence score above a threshold of 0.7 were retained for further analysis, ensuring that only the highest-confidence structures were considered as potential candidates for downstream interpretation.

### Docking experiments and analysis

We applied docking simulations to predict the interaction between human GR-LBD (PDB = 7PRW) and human NIPBL (PDB = 6WG3) using the pyDock docking and scoring method (39). First, protein molecules were prepared by removing all cofactors and heteroatoms, and missing side chains were modeled with SCWRL 3.0 (55). The Fast Fourier Transform (FFT)-based docking program FTDock (56) (with electrostatics and 0.7 Å grid resolution) was used to generate 10,000 rigid-body docking poses. Then, pyDock scoring was used based on energy terms previously optimized for rigid-body docking (39). The pyDock binding energy is composed of accessible surface area-based desolvation, Coulombic electrostatics, and Van der Waals (VdW) energy terms. Electrostatics and VdW contributions were limited to -1.0/+1.0 and 1.0 kcal/mol for each inter-atomic energy value, respectively, to avoid excessive penalization from possible clashes derived from the rigid-body approach.

### RNA preparation

Two biological replicates of 3134 cl-15 cells expressing different NIPBL constructs were sub-cultured in medium containing 10% charcoal-stripped FBS for 3 days with or without 1 mM of IPTG as indicated. Samples were treated with 100 nM of Dex or equivalent 0.1% of EtOH for 2 hours, then RNA was processed for extraction following the manufacturer’s protocol (Nucleospin RNA extraction kit, Macherey-Nagel) and stored at -80°C.

### RNA-sequencing

RNA libraries were constructed according to the Illumina Stranded Total RNA Prep, Ligation with Ribo-Zero Plus. Samples were pooled, and paired-end sequencing was performed on NovaSeq 6000 S2. Raw reads were demultiplexed using Illumina Bcl2fastq v2.20. Reads were trimmed for adapters and low-quality bases using Cutadapt1.18 before alignment with the reference genome (mm10) and the annotated transcripts (GENCODE M21) using STAR 2.7.0f. Raw tag counts of exon regions at the gene level were obtained using featureCounts in subread 2.0.3. Default DESeq2 size factors were used to normalize the data. The significance of gene expression changes was evaluated based on Wald-statistics (FDR<0.05) and log_2_fold-change (>log_2_(1.5), <-log_2_(1.5)) using DESeq2 library (1.42.1). The EtOH and Dex-treated RNA-seq samples were sequenced separately. To account for the variability in the baseline gene expression between the two sequencing batches, paired analysis was performed to evaluate differential gene expression. For clustered heatmaps, the raw tag count data was variance stabilizing transformed using VST function in DESeq2, and the z-score of the transformed data was visualized as clustered heatmaps using pheatmap R library (1.0.12) function with hierarchical, euclidean, ward.D2 methods.

