## Supplementary figures and tables for "Transcription factors form a ternary complex with NIPBL/MAU2 to localize cohesin at enhancers"

**This PDF includes**

Figures S1 to S4

Tables S1 and S2

Videos S1 and S2

**Figure S1**

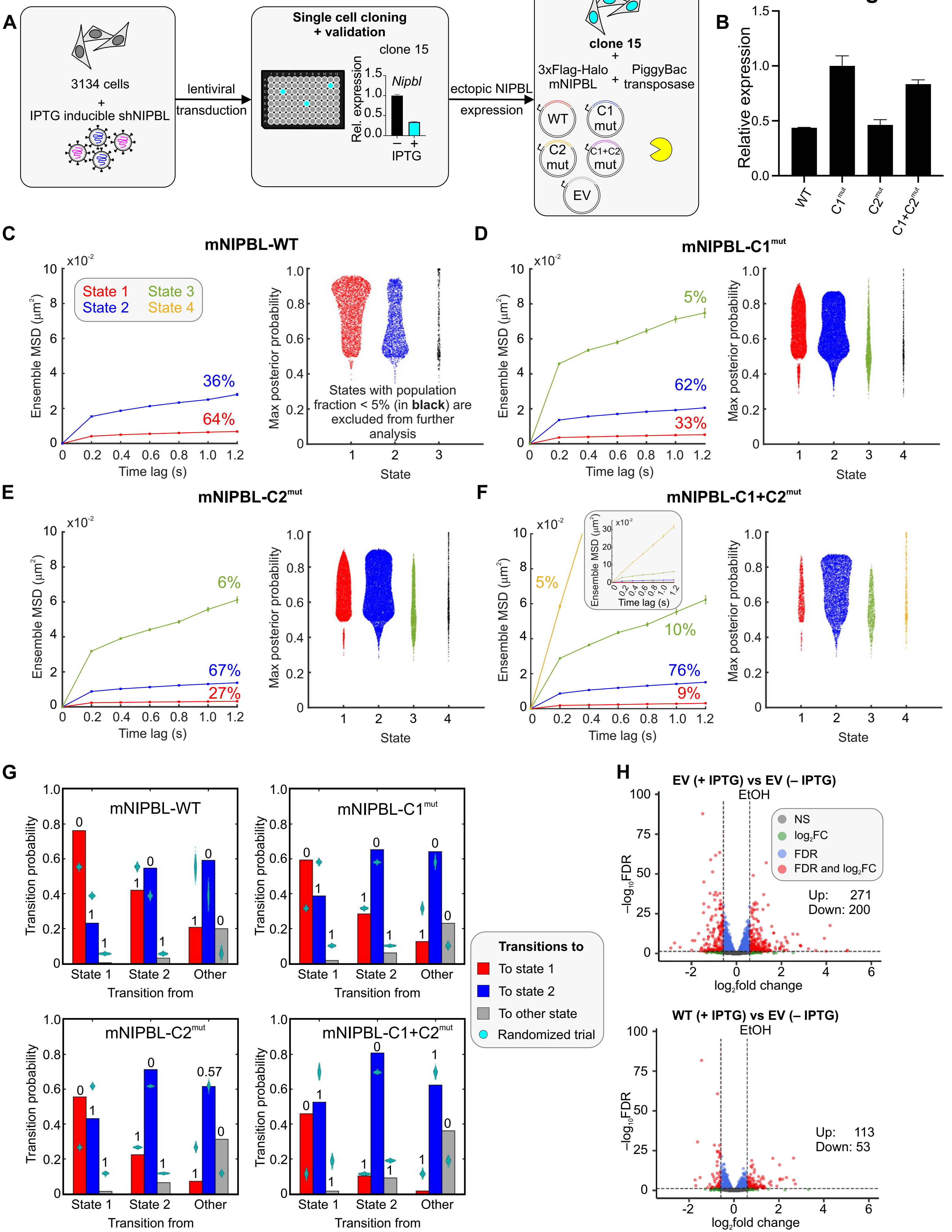
A
3134 cells
+
IPTG inducible shNIPBL
lentiviral transduction
Single cell cloning + validation
clone 15
Nipbl
Rel. expression
IPTG
ectopic NIPBL expression
clone 15
+
3xFlag-Halo mNIPBL
+
PiggyBac transposase
WT
C1 mut
C2 mut
C1+C2 mut
EV
B
Relative expression
WT
C1mut
C2mut
C1+C2mut
C
mNIPBL-WT
Ensemble MSD (μm²)
x10-2
State 1
State 2
State 3
State 4
36%
64%
Time lag (s)
Max posterior probability
States with population fraction < 5% (in black) are excluded from further analysis
State
D
mNIPBL-C1mut
5%
62%
33%
E
mNIPBL-C2mut
6%
67%
27%
F
mNIPBL-C1+C2mut
5%
10%
76%
9%
G
Transition probability
Transition from
State 1
State 2
Other
mNIPBL-WT
mNIPBL-C1mut
mNIPBL-C2mut
mNIPBL-C1+C2mut
Transitions to
To state 1
To state 2
To other state
Randomized trial
H
EV (+ IPTG) vs EV (– IPTG)
EtOH
NS
log2FC
FDR
FDR and log2FC
Up: 271
Down: 200
−log10FDR
log2fold change
WT (+ IPTG) vs EV (– IPTG)
EtOH
Up: 113
Down: 53

**Figure S1: The role of Leu-rich clusters in NIPBL dynamics and biology**

(A) Schematic describing the generation of the IPTG-inducible NIPBL-KD cell line, and subsequent ectopic expression of mNIPBL-WT, $C1^{mut}$, $C2^{mut}$, $C1+C2^{mut}$, or empty vector (EV, 3xFLAG-Halo alone). The middle panel shows the reduction in *Nipbl* mRNA upon 72 h of 1 mM IPTG treatment.

(B) RNA-seq reads of the ectopic mNIPBL constructs normalized to the highest expression. Error bars represent the standard deviation.

(C–F) (Left) Ensemble mean-squared displacement (MSD) curves for the indicated species. State 1 is depicted in red, state 2 in blue, state 3 in green, and state 4 in yellow. The numbers above the MSD curves indicate the population fraction of the respective states. Error bars denote the standard error. (Right) Swarmcharts of the maximum posterior probability for each state detected by pEMv2. Colors match those of the states presented in the MSD plots. States with less than 5% population fraction are presented in black. Inset of panel F (left) shows the same MSD curve as panel F with higher y-axis limits.

(G) Transition probability bar plots for transitions from the states indicated on the x-axes to state 1 (red), state 2 (blue), or any 'other' state (grey). The cyan swarmcharts show the transition probabilities calculated from 1000 randomized ensembles of trajectories and the numbers above the bars denote the fraction of randomized ensembles that have a transition probability higher than the respective calculated transition probability.

(H) Volcano plots of the $-\log_{10}$FDR and $\log_2$(fold change) showing the differential expression of genes under basal conditions (cells grown in charcoal-stripped FBS containing medium). (Top) cl15-EV (+ IPTG) vs cl-15-EV (without IPTG). (Bottom) cl15-WT (+ IPTG) vs cl15-EV (without IPTG). The black circles denote genes that do not meet either the FDR or fold-change cut-offs, green circles denote those that only meet the fold-change cut-off, blue circles denote those that only meet the FDR cut-off, and the red circles denote the genes that meet both the fold-change and FDR cut-offs. Dashed lines indicate the FDR and fold-change thresholds. (Insets) the number of up and down-regulated genes in each condition.

Related to Figure 1.

**Figure S2**

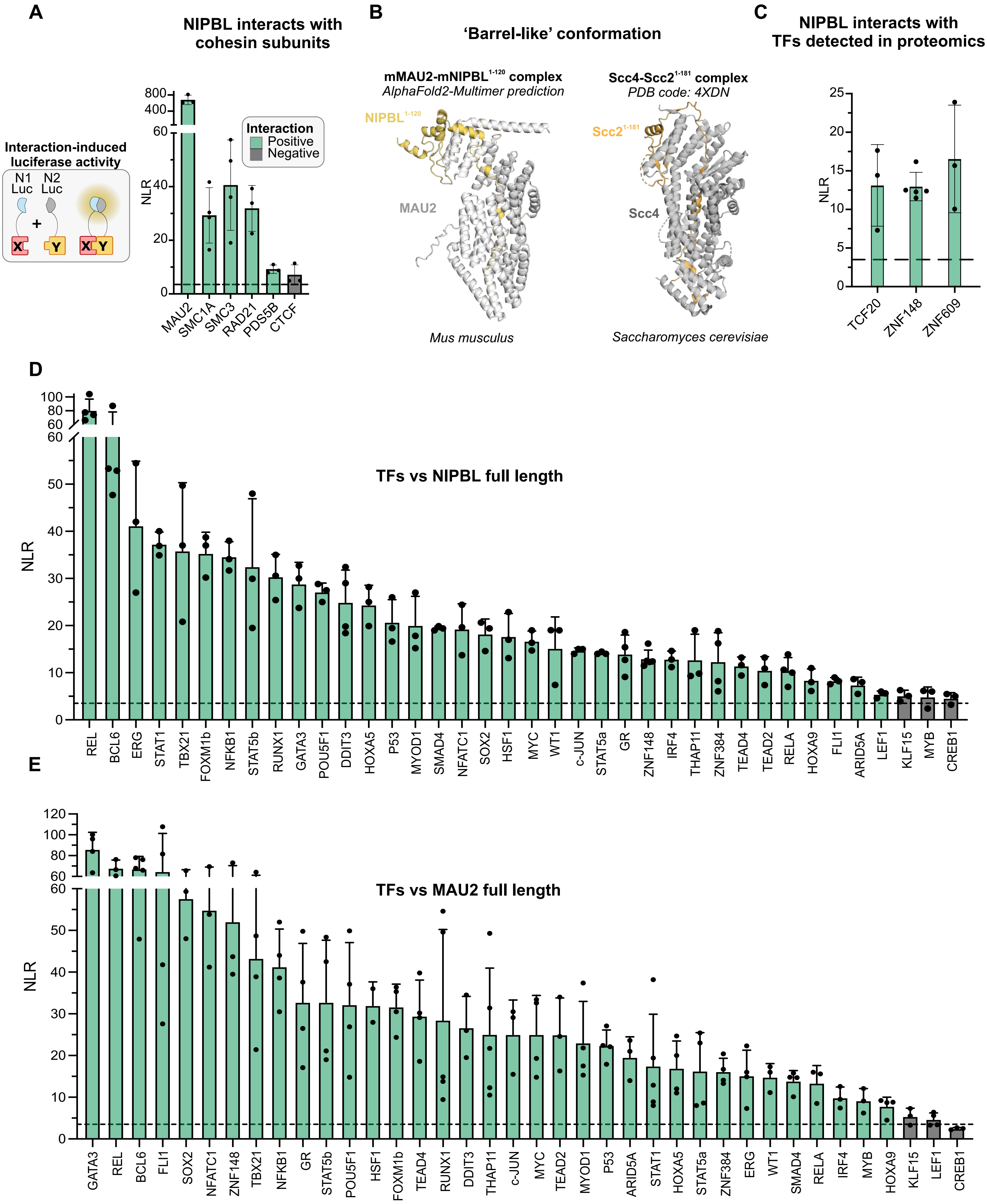
A
NIPBL interacts with cohesin subunits
Interaction-induced luciferase activity
N1 Luc
N2 Luc
X
Y
Interaction
Positive
Negative
NLR
MAU2
SMC1A
SMC3
RAD21
PDS5B
CTCF
B
'Barrel-like' conformation
mMAU2-mNIPBL1-120 complex
AlphaFold2-Multimer prediction
NIPBL1-120
MAU2
Mus musculus
Scc4-Scc21-181 complex
PDB code: 4XDN
Scc21-181
Scc4
Saccharomyces cerevisiae
C
NIPBL interacts with TFs detected in proteomics
NLR
TCF20
ZNF148
ZNF609
D
TFs vs NIPBL full length
NLR
REL
BCL6
ERG
STAT1
TBX21
FOXM1b
NFKB1
STAT5b
RUNX1
GATA3
POU5F1
DDIT3
HOXA5
P53
MYOD1
SMAD4
NFATC1
SOX2
HSF1
MYC
WT1
c-JUN
STAT5a
GR
ZNF148
IRF4
THAP11
ZNF384
TEAD4
TEAD2
RELA
HOXA9
FLI1
ARID5A
LEF1
KLF15
MYB
CREB1
E
TFs vs MAU2 full length
NLR
GATA3
REL
BCL6
FLI1
SOX2
NFATC1
ZNF148
TBX21
NFKB1
GR
STAT5b
POU5F1
HSF1
FOXM1b
TEAD4
RUNX1
DDIT3
THAP11
c-JUN
MYC
TEAD2
MYOD1
P53
ARID5A
STAT1
HOXA5
STAT5a
ZNF384
ERG
WT1
SMAD4
RELA
IRF4
MYB
HOXA9
KLF15
LEF1
CREB1

**Figure S2: Gaussia protein-fragment complementation assay (gPCA) detects direct interactions between proteins**

(A) (Left) Schematic representation of a gPCA experiment. (Right) Normalized luminescence ratio (NLR) for gPCA experiments measuring interactions between hNIPBL-WT and cohesin sub-units and CTCF.

(B) Comparison of the mouse MAU2-NIPBL complex prediction (left) with the crystal structure of the yeast Scc4-Scc2 complex (PDB: 4XDN) (right). Notice the structural resemblance of the 'barrel-like' conformation of MAU2/Scc4 surrounding the NIPBL/Scc2 region.

(C–E) NLRs for gPCA experiments between (C) NIPBL-WT and three transcription factors (TFs) detected in the proteomics experiments, (D) NIPBL-WT and the indicated TFs, (E) hMAU2 and the indicated TFs.

Positive interactions are indicated in green while negative interactions are in black. Error bars show the standard deviation across multiple measurements.

Related to Figure 2.

**Figure S3**

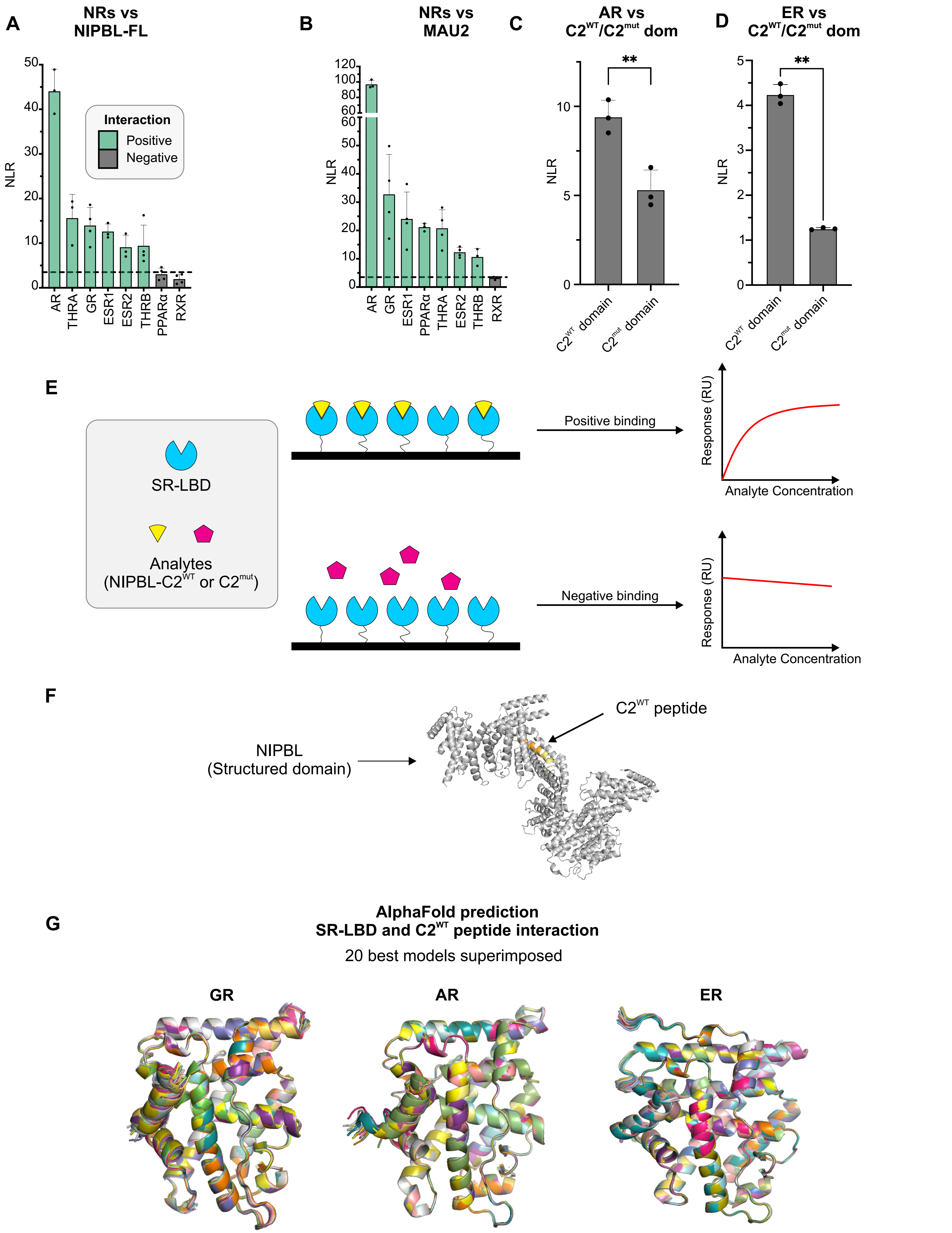
A
NRs vs NIPBL-FL
NLR
Interaction
Positive
Negative
AR
THRA
GR
ESR1
ESR2
THRB
PPARα
RXR
B
NRs vs MAU2
NLR
AR
GR
ESR1
PPARα
THRA
ESR2
THRB
RXR
C
AR vs C2^WT/C2^mut dom
NLR
**
C2^WT domain
C2^mut domain
D
ER vs C2^WT/C2^mut dom
NLR
**
C2^WT domain
C2^mut domain
E
SR-LBD
Analytes
(NIPBL-C2^WT or C2^mut)
Positive binding
Negative binding
Response (RU)
Analyte Concentration
F
NIPBL
(Structured domain)
C2^WT peptide
G
AlphaFold prediction
SR-LBD and C2^WT peptide interaction
20 best models superimposed
GR
AR
ER

**Figure S3: NIPBL-C2 interacts with steroid receptor ligand-binding domains**

(A–B) NLRs of interactions measured by gPCA between nuclear receptors and (A) full-length NIPBL-WT and (B) MAU2. Positive interactions are indicated in green while negative interactions are in black. Error bars represent the standard deviation.

(C-D) NLRs of gPCA interactions between NIPBL-$C2^{WT}$ and $C2^{mut}$ domains and (C) AR and (D) ER. **$p<0.01$ (paired t-test).

(E) Schematic of a surface plasmon resonance (SPR) experiment.

(F) AlphaFold prediction of the structure of the structured region of human NIPBL containing the C2 LxxLL motif cluster. Residues from the synthesized peptide used for the SPR experiment are depicted in yellow and the exact LxxLL motif sequence used for AlphaFold2-Multimer predictions is colored in ochre.

(G) Superposition of the 20 best AlphaFold2-Multimer models showing the interaction between the $C2^{WT}$ peptide and (left) human GR, (middle) human AR, and (right) human ER.

Related to Figure 3.

**Figure S4**

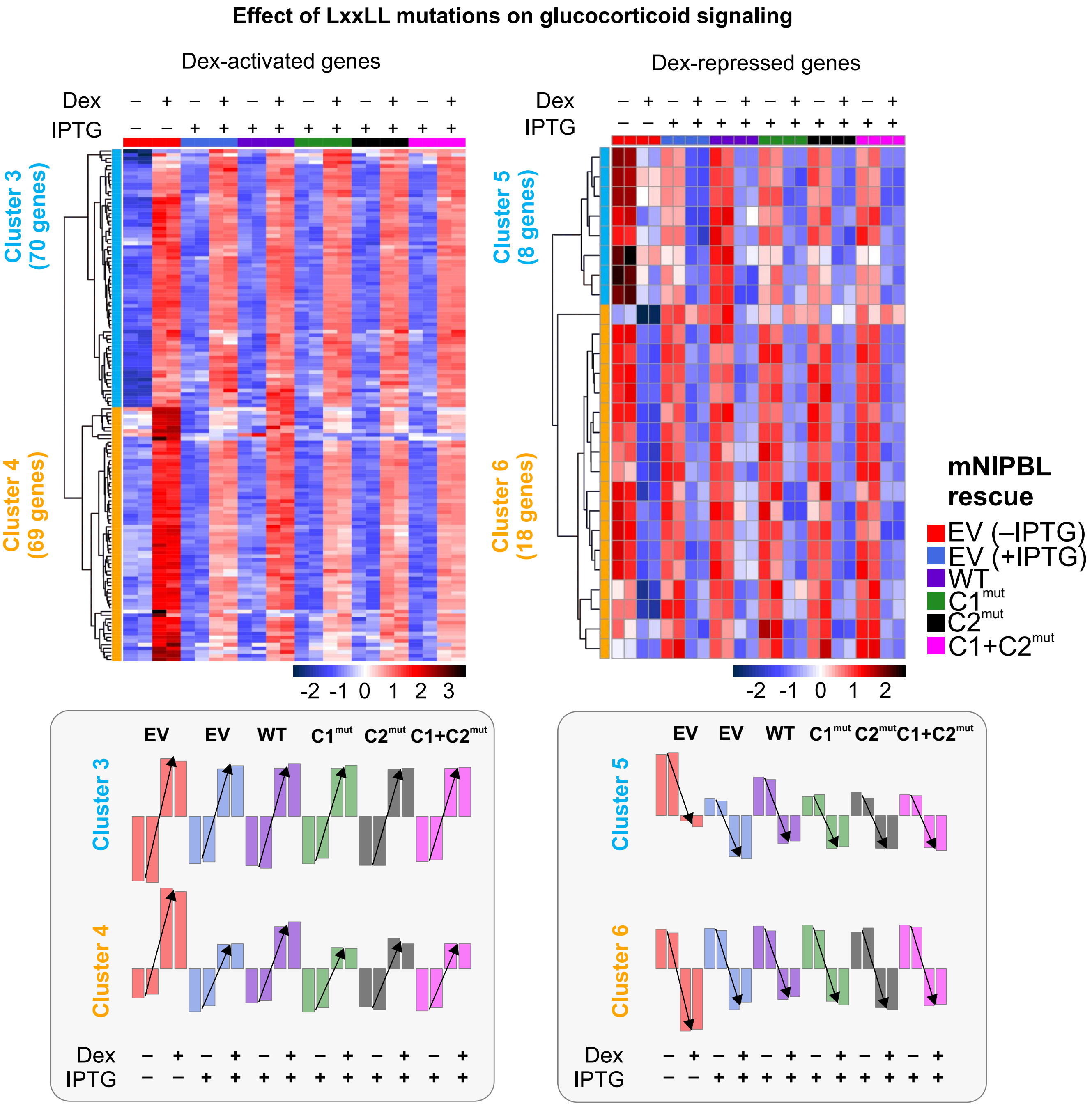
Effect of LxxLL mutations on glucocorticoid signaling
Dex-activated genes
Dex-repressed genes
Dex
IPTG
Cluster 3 (70 genes)
Cluster 4 (69 genes)
Cluster 5 (8 genes)
Cluster 6 (18 genes)
-2 -1 0 1 2 3
-2 -1 0 1 2
mNIPBL rescue
EV (–IPTG)
EV (+IPTG)
WT
C1mut
C2mut
C1+C2mut
EV EV WT C1mut C2mut C1+C2mut
Cluster 3
Cluster 4
Cluster 5
Cluster 6

**Figure S4: NIPBL mutants alter a subset of glucocorticoid receptor-regulated genes**

Heatmap of genes whose Dex-induced fold change is significantly affected (FDR = 0.1) in at least one of the NIPBL-KD and/or NIPBL-WT/C1$^{mut}$/C2$^{mut}$/C1+C2$^{mut}$ conditions relative to EV (no IPTG, with endogenous NIPBL), separated by those activated (left) or repressed (right) in response to Dex treatment. The bottom bar plots indicate the average expression of the genes in the indicated clusters.

Related to Figure 4.

**Table S1:** Sources for the ORFs used for the gPCA experiments

| **ORF** | **SOURCE** |
|---|---|
| ARID5A | Human orfeome database v7.1 |
| BCL6 | Human orfeome database v7.1 |
| CREB1 | Human orfeome database v7.1 |
| E2F1 | Human orfeome database v7.1 |
| ESR1 | Human orfeome database v7.1 |
| ESR2 | Human orfeome database v7.1 |
| FOXM1b | Human orfeome database v7.1 |
| GATA3 | Human orfeome database v7.1 |
| HOXA5 | Human orfeome database v7.1 |
| HOXA9 | Human orfeome database v7.1 |
| HSF1 | Human orfeome database v7.1 |
| JUN | Human orfeome database v7.1 |
| LEF1 | Human orfeome database v7.1 |
| MYC | Human orfeome database v8.1 |
| MYB | Human orfeome database v7.1 |
| MYOD1 | Human orfeome database v7.1 |
| NFKB1 | Yvette Habraken Lab |
| POU2F1 | Human orfeome database v7.1 |
| REL | Human orfeome database v7.1 |
| RELA | Human orfeome database v7.1 |
| RUNX1 | Human orfeome database v7.1 |
| SMAD4 | Human orfeome database v8.1 |
| STAT1 | Human orfeome database v7.1 |
| STAT5A | Human orfeome database v7.1 |
| STAT5B | Human orfeome database v7.1 |
| TBX21 | Human orfeome database v7.1 |
| THAP11 | Human orfeome database v7.1 |
| THRA | Human orfeome database v7.1 |
| TP53 | Human orfeome database v7.1 |
| WT1 | Human orfeome database v7.1 |
| NIPBL | This study |

| | |
|---|---|
| MAU2 | This study |
| SMC1A | Addgene #32363 |
| SMC3 | Addgene #156447 |
| RAD21 | Addgene #156445 |
| PDS5B | Addgene #156442 |
| CTCF | Addgene #40801 |
| AR | Gordon Hager Lab |
| THRB | Gordon Hager Lab |
| GR | Gordon Hager Lab |
| PPAR$\alpha$ | Gordon Hager Lab |
| RXR | Gordon Hager Lab |
| DDIT3 | Human orfeome database v8.1 |
| NFATC1 | Human orfeome database v8.1 |
| SOX2 | Human orfeome database v8.1 |
| c-JUN | Yvette Habraken Lab |
| ZNF148 | Human orfeome database v8.1 |
| IRF4 | Human orfeome database v8.1 |
| ZNF384 | Human orfeome database v7.1 |
| TEAD2 | Human orfeome database v8.1 |
| TEAD4 | Human orfeome database v8.1 |
| FLI1 | Human orfeome database v7.1 |
| KLF15 | Human orfeome database v8.1 |
| TCF20 | PMID 35074918 |
| ZNF609 | PMID 28041881 |
| GR-$\Delta$NTD | Cloned from pDONOR-GR |
| GR-$\Delta$LBD | Cloned from pDONOR-GR |
| NIPBL C1$^{WT}$ domain | Cloned from pPB-3xFLAG-Halo-NIPBL-WT |
| NIPBL C2$^{WT}$ domain | Cloned from pPB-3xFLAG-Halo-NIPBL-WT |
| NIPBL C2$^{mut}$ domain | Cloned from pPB-3xFLAG-Halo-NIPBL-C2$^{mut}$ |

**Table S2:** Forward and reverse primers for the cloning of gPCA ORFs

| | **PRIMER FWD AttB1** | **PRIMER REV AttB2** |
|---|---|---|
| NIPBL C1$^{WT}$ domain | GGGGACAACTTTGTACAAAAAAGTTGGCATGGCCGGAATCGCTTCTCTGAC | GGGGACAACTTTGTACAAGAAAGTTGCATGCCGCTCTGCACGTATC |
| NIPBL C2$^{WT}$ domain | GGGGACAACTTTGTACAAAAAAGTTGGCATGTACATCCAGATGGTTACAGCTCTGG | GGGGACAACTTTGTACAAGAAAGTTGCTCTCCCAGGACTCTCAGGATC |
| NIPBL C2$^{mut}$ domain | GGGGACAACTTTGTACAAAAAAGTTGGCATGTACATCCAGATGGTTACAGCTCTGG | GGGGACAACTTTGTACAAGAAAGTTGCTCTCCCAGGACTCTCAGGATC |
| GR-ΔNTD | GGGGACAACTTTGTACAAAAAAGTTGGCATGAGACCAGATGTGAGTTCTCCTCC | GGGGACAACTTTGTACAAGAAAGTTGTTTCTGATGAAACAGAAGCTTTTTGATATTTCCATTTG |
| GR-ΔLBD | GGGGACAACTTTGTACAAAAAAGTTGGCATGGACTCCAAAGAATCCTTAGCTCCCC | GGGGACAACTTTGTACAAGAAAGTTGGTCTTGTGAGACTCCTGCAGTG |
| NIPBL | GGGGACAACTTTGTACAAAAAAGTTGGCATGAACGGCGACATGCCTCACGTGC | GGGGACAACTTTGTACAAGAAAGTTGAAGAGCTTGTGCCGTCCTTAGCAGCG |
| MAU2 | GGGGACAACTTTGTACAAAAAAGTTGGCATGGCGGCACAGGCGGCGG | GGGGACAACTTTGTACAAGAAAGTTGCAGAAGGCTGGCCAGGCTGG |
| SMC1A | GGGGACAACTTTGTACAAAAAAGTTGGCATGGGGTTCCTGAAACTGATTGAG | GGGGACAACTTTGTACAAGAAAGTTGCTGCTCATTGGGGTTGGGG |
| SMC3 | GGGGACAACTTTGTACAAAAAAGTTGGCATGTACATCAAGCAGGTGATCATCCAG | GGGGACAACTTTGTACAAGAAAGTTGACCATGCGTGGTATCGTCTTC |
| RAD21 | GGGGACAACTTTGTACAAAAAAGTTGGCATGTTCTACGCACATTTTGTCCTCAG | GGGGACAACTTTGTACAAGAAAGTTGGATAATATGGAACCGTGGTCCAGGG |
| PDS5B | GGGGACAACTTTGTACAAAAAAGTTGGCATGGCTCATTCAAAGACAAGGACC | GGGGACAACTTTGTACAAGAAAGTTGTCGTCTCTCTCGTTTGGAGCTTC |
| CTCF | GGGGACAACTTTGTACAAAAAAGTTGGCATGGAAGGTGAGGCGGTTG | GGGGACAACTTTGTACAAGAAAGTTGCCGGTCCATCATGCTGAGG |
| AR | GGGGACAACTTTGTACAAAAAAGTTGGCATGGAAGTGCAGTTAGGGCTGGG | GGGGACAACTTTGTACAAGAAAGTTGCTGGGTGTGGAAATAGATGGGC |
| THRB | GGGGACAACTTTGTACAAAAAAGTTGGCATGGCGATCGCCATGACTCCC | GGGGACAACTTTGTACAAGAAAGTTGAACATCCTCGAACACTTCCAAGAAC |

| | | |
|---|---|---|
| GR | GGGGACAACTTTGTACAAAAAAGTTGGCATGGACTCCAAAGAATCCTTAGCTCCC | GGGGACAACTTTGTACAAGAAAGTTGTTTCTGATGAAACAGAAGCTTTTTGATATTTCC |
| PPARα | GGGGACAACTTTGTACAAAAAAGTTGGCATGGTGGACACGGAAAGCCCAC | GGGGACAACTTTGTACAAGAAAGTTGAACGTACATGTCCCTGTAGATCTCC |
| RXR | GGGGACAACTTTGTACAAAAAAGTTGGCATGGACACCAAACATTTCCTGCCG | GGGGACAACTTTGTACAAGAAAGTTGAACAGTCATTTGGTGCGGC |
| TCF20 | GGGGACAACTTTGTACAAAAAAGTTGGCATGCAGTCCTTTCGGGAGCAAAGC | GGGGACAACTTTGTACAAGAAAGTTGTCTCCACAGTCTCACCTTGTGCTTAG |
| ZNF609 | GGGGACAACTTTGTACAAAAAAGTTGGCATGTCCTTGAGCAGTGGAGCCTG | GGGGACAACTTTGTACAAGAAAGTTGCCTCCGGGGGGGTGG |

**Video S1: Representative fast SMT movies for NIPBL-WT and mutants**

Montage of representative SMT movies collected with the fast SMT protocol (exposure time = 12 ms, interval = 12 ms). Left to right: mNIPBL-WT, $C1^{mut}$, $C2^{mut}$, and $C1+C2^{mut}$. Trajectories longer than 7 frames are shown in red. Scale bar = 2 μm and playback speed in 125 frames/s.

Related to Figure 1.

**Video S2: Representative intermediate SMT movies for NIPBL-WT and mutants**

Montage of representative SMT movies collected with the intermediate SMT protocol (exposure time = 10 ms, interval = 200 ms). Left to right: mNIPBL-WT, $C1^{mut}$, $C2^{mut}$, and $C1+C2^{mut}$. Trajectories longer than 7 frames are shown in red. Scale bar = 2 μm and playback speed in 50 frames/s.

Related to Figure 1.
